# Attention Decoding at the Cocktail Party: Preserved in Hearing Aid Users, Reduced in Cochlear Implant Users

**DOI:** 10.64898/2025.12.22.695344

**Authors:** Constantin Jehn, Jasmin Riegel, Tobias Reichenbach, Anja Hahne, Niki Katerina Vavatzanidis

## Abstract

Users of hearing aids (HAs) and cochlear implants (CIs) experience significant difficulty understanding a target speaker in multi-talker environments or when other background noise is present. Segregation of a particular voice from background noise occurs partly through enhanced cortical tracking of amplitude fluctuations in the target signal. Measuring a person’s cortical tracking allows decoding their focus of attention and may be used for neurofeedback in hearing devices, potentially aiding their users with speech-in-noise comprehension. Most studies on cortical speech tracking have employed typical hearing (TH) individuals, whereas studies in people with hearing impairment whose cortical tracking may differ are still scarce. The objective of this study was to compare cortical speech tracking of HA (n=29) and CI users (n=24) to that of age-matched TH individuals (n=29). We recorded EEG data while the participants attended one of two competing talkers (one with a female and one with a male voice), in a free-field acoustic environment. Importantly, HA users as well as CI users used their personal, clinically-fitted devices. Cortical speech tracking was assessed through linear backward and forward models that related the EEG data to the speech envelope. For the CI users, electrical artifacts stemming from the implant were addressed through a bespoke method for artifact rejection. We found that the HA group exhibited cortical tracking and attentional modulation that were largely comparable to those of the TH group. CI users also showed successful cortical tracking. However, they displayed a profound deficit in attentional modulation, seen in the significantly poorer neural segregation of the attended vs. the ignored speech streams. These results shed light on a neurobiological mechanism for speech-in-noise comprehension and have implications for neurofeedback in hearing devices.

## 1 Introduction

One in five people globally (1.57 billion) were affected by hearing loss in 2019, and 430 million of them had moderate to severe hearing loss (*≥* 35 dB without adjustment). With ageing and population growth, these numbers are projected to increase to 2.45 billion and 698 million by 2050 (Haile et al., 2021). Untreated hearing loss has been associated with cognitive decline and diminished social wellbeing as it impairs communication and social participation (Yeo et al., 2023). Moderate to severe hearing loss can be treated using hearing aids (HAs), which restore audibility by amplifying incoming acoustic signals (Hoppe and Hesse, 2017). In cases of severe to profound sensorineural hearing loss, cochlear implants (CIs) provide an effective intervention by bypassing the auditory periphery and directly stimulating the auditory nerve via an implanted electrode array (Gaylor et al., 2013). Although one million CIs have been implanted by 2022, thus improving the lives of many severely hearing impaired individuals, the perception of pitch and temporal fine structure (TFS) is still poor for CI users compared to typical hearing (TH) individuals or users of hearing aids (Wouters et al., 2015; Zeng, 2022). As a result, many CI users face substantial difficulties understanding speech in the presence of background noise, which is required in many social situations or at work (Fowler et al., 2021). By contrast, hearing aid users perceive sound through amplification and otherwise natural mechanotransduction in the inner ear. They thus have better pitch and TFS perception compared to CI users (Looi et al., 2008). However, they also experience greater difficulty and increased listening effort when communicating in multi-talker, so-called cocktail-party environments, compared to TH individuals (Harkins and Tucker, 2007; Marrone et al., 2008).

When we listen to speech, the cortical activity in the low-frequency range of 1*−*12 Hz tracks the amplitude fluctuations in the speech signal (Ding and Simon, 2014; Giraud and Poeppel, 2012). This cortical speech tracking plays an important role in segregating a speech signal from background noise, such as another talker: the cortical tracking of an attended speaker is stronger and occurs at a different phase than that of an ignored speech stream (Ding and Simon, 2012; Horton et al., 2013; Mesgarani and Chang, 2012). The difference in the cortical tracking of an attended and an ignored speech stream is so pronounced in typically hearing individuals that it allows for the decoding of auditory attention from a few seconds of non-invasive electroencephalographic (EEG) recordings (Geirnaert et al., 2021; O’Sullivan et al., 2015). This form of auditory attention decoding could function as a neurofeedback loop in hearing devices by identifying the attended speaker and enhancing their speech stream, thereby reducing background noise and supporting the user in speech-in-noise comprehension (Hjortkjær et al., 2025).

Against this background, as multi-talker scenarios continue to pose a major challenge for users of assistive hearing technologies, there is growing interest in investigating cortical speech tracking and auditory attention decoding in individuals with hearing impairment. Therefore, the influence of peripheral hearing loss on cortical speech tracking has been examined in several studies using selective attention paradigms. In these studies, cortical speech tracking has been typically quantified using encoding, decoding models, or a combination of both. Encoding approaches, often implemented via temporal response functions (TRFs), model how features of the speech signal predict neural responses, thereby characterizing stimulus–brain representations across time. By contrast, decoding approaches reconstruct speech features or infer attentional focus directly from neural activity.

Using these approaches, Fuglsang et al. found that hearing-impaired individuals demonstrated higher encoding and decoding accuracy than an age-matched control group, suggesting enhanced neural tracking of attended speech (Fuglsang et al., 2020). Importantly, participants with hearing loss did not use their regular hearing aids during the experiment; instead, audibility was compensated for by applying a frequency-specific gain to the stimuli. Extending this line of research, Decruy et al. observed increased neural envelope tracking of the target speaker in hearing-impaired participants compared to TH controls during a speech-on-speech task (Decruy et al., 2020). These findings suggest that hearing loss may be associated with compensatory enhancements of speech tracking at the cortical level under complex listening conditions. However, an earlier study reported a reduced differential in neural tracking between attended and ignored speech streams in hearing-impaired individuals (Petersen et al., 2017), which contrasts with the enhanced tracking observed in more recent studies. In their study, participants used hearing aids with quasi-linear amplification and the stimuli were streamed directly to the devices. An open question remains as to how neural tracking manifests in hearing aid users while they are actively wearing their devices in a more realistic free-field listening environment.

For CI users, studies by Nogueira et al. showed that cortical envelope tracking can be used to decode selective attention for monaural and dichotic stimuli presentation in this population as well (Nogueira and Dolhopiatenko, 2022; Nogueira et al., 2019), despite severe electrical artifacts in the EEG caused by the CIs (Wagner et al., 2018). Following these findings, Paul et al. investigated at which latencies the cortical differentiation of speakers appears in CI users, employing a temporal response function (TRF) analysis (Paul et al., 2020). They found that cortical talker representation is quite similar and only varies for later peaks: 250 ms in CI users compared to 150 ms in typical hearing individuals.

While previous research has investigated cortical speech tracking and attentional modulation in hearing-impaired listeners, a direct comparison of CI users, HA users, and an age-matched typically hearing (TH) control group within the same selective attention paradigm has been missing. Moreover, many of the cited studies presented stimuli directly via cable to CIs or employed simplified amplification strategies, leaving unanswered the question of how modern, clinically-fitted devices impact neural processing in realistic listening environments.

The present study directly addresses these gaps. We investigate cortical speech tracking in these three distinct, age-matched groups in both single-talker and competing-talker conditions. By using a free-field setup and the participants’ own clinically fitted devices, we achieve the high ecological validity necessary to assess this question. This work, therefore, provides the first comprehensive, side-by-side characterization of cortical speech tracking in individuals utilizing both forms of hearing assistive technology.

## 2 Methods

### 2.1 Participants

In total, 95 participants were recruited. All participants provided written informed consent to take part in the study. The experiment was approved by the science ethics committee of Technische Universität Dresden (protocol SR+BO-EK-47022024) on May 21, 2024. All procedures were conducted in accordance with the Declaration of Helsinki.

#### Cochlear implant group (CI)

The first group comprised 30 bilaterally implanted CI users. Data sets of six participants were excluded: Four data sets suffered from technical issues during the recording session, and two participants were unable to finish the study. Eventually, data from 24 bilateral CI users were analyzed (CI experience: median = 10 years, range = 2*−*28 years). Due to the demanding nature of the selective attention paradigm in this study, we only included CI users with relatively high levels of speech comprehension. Specifically, participants were required to score at least 60% on the Freiburg monosyllabic word recognition test (Freiburger Einsilber) in quiet measured at 65 dB for their better-performing ear (each ear was tested separately) (Hahlbrock, 1970). We furthermore obtained speech comprehension scores in noise at the sentence level (65 dB signal, 60 dB noise, free-field environment with both CI sides) with the Hochmair–Schulz–Moser (HSM) sentence test (Hochmair-Desoyer et al., 1997). The implantation history of the CI group was largely post-lingual (n=20), with one pre-lingual and three peri-lingual (implanted before/during speech acquisition) participants. To control for this potential confound, we compared the post-lingual (n=20) and combined pre/peri-lingual (n=4) groups on our key speech-on-speech performance metrics: decoding accuracy, comprehension scores, and listening effort (all in the competing speaker condition). Further demographic information of the 24 included CI users is summarized in Table 1. Participant-level demographic and technical details are provided in Appendix B (Table 4).

**Table 1:** Demographics of the 24 studied bilateral CI users.

| Metric | Age<br>(years) | Age at fit<br>(years) | CI experience<br>(years) | Monosyllable<br>65 dB (%)<sup>1</sup> | Monosyllable<br>80 dB (%)<sup>1</sup> | HSM<br>in noise (%)<sup>2</sup> | Sex |
| --- | --- | --- | --- | --- | --- | --- | --- |
| Median | 56 | 43 | 10 | 85 | 90 | 23.5 | 14F 10M |
| Range | 25 - 81 | 2 - 74 | 2 - 28 | 60 - 100 | 75 - 100 | 0 - 82 | — |
| Std. | 17.1 | 19.6 | 6.7 | 8.1 | 6.3 | 22.8 | — |

<sup>1</sup> Monaural Freiburg monosyllabic word test in quiet at 65 dB and 80 dB, scored for the better ear.
<sup>2</sup> Binaural Hochmair–Schulz–Moser (HSM) sentence test in noise (65 dB signal, 60 dB noise).

#### Hearing aid group (HA)

Our HA cohort comprised 29 bilateral HA users. They completed both the Freiburg monosyllabic word test in quiet measured at 65 dB and the HSM test (65 dB signal, 60 dB noise), with their hearing aids on, to enable a performance comparison with the CI group. Both tests were conducted in a free-field environment. However, the word test for five participants was administered via headphones. As this method is non-compliant with standard protocols for HA users (Hamzavi et al., 2001), word test data from these individuals were excluded from the analysis. All other HA users completed the monaural word test with the contralateral ear occluded using an earbud and an additional over-ear hearing protection.

In addition, unaided pure-tone audiometry scores were obtained either from their hearing-care professionals or tested before the study. The time between the audiometry measurement and the study was less than 2 years for 27 participants (median: 8 months, range: 0 *−* 21 months, outliers: 64 and 85 months). During the experiment, they used the everyday setting of their HAs, most of which we expect to be adaptive, so cortical tracking could be assessed during the use of their own assistive hearing technology.

#### Typically hearing group (TH)

Furthermore, 33 individuals with typical hearing were recruited as an age-matched control group. The inclusion criterion required an audiometric threshold better than 25 dB for each ear, which was validated through pure-tone audiometry (PTA4) before the EEG experiment. Four participants did not meet this criterion and were thus excluded from the analysis, resulting in a final control group of 29 individuals.

#### Distributions of sex, age and hearing performance across groups

Fig. 1 summarizes the demographic and audiometric profiles of the three groups. In panel **A**, we consistently see slightly more female participants in all groups. A chi-squared test of independence confirmed that sex distribution did not differ between groups (*χ*^2^ (2) = 0.100, *p* = 0.951). The corresponding effect size was negligible (Cramér’s V = 0.034), indicating no meaningful association between sex and group. Panel **B** shows a broad age distribution across all cohorts, ranging from 19 to 81 years. The inclusion criteria for the CI group, including bilateral implantation and a relatively high level of speech comprehension, ensured that, despite the broad age range, participants were comparable. We confirmed that age distributions did not differ between the three cohorts (*H*(2) = 1.03*, p* = 0.597), by applying the non-parametric Kruskal–Wallis test, as the age distributions of the TH and CI groups violated normality. The effect size was negligible 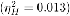, indicating that group membership explained only 1.3% of the variance in age.

**Fig. 1.**
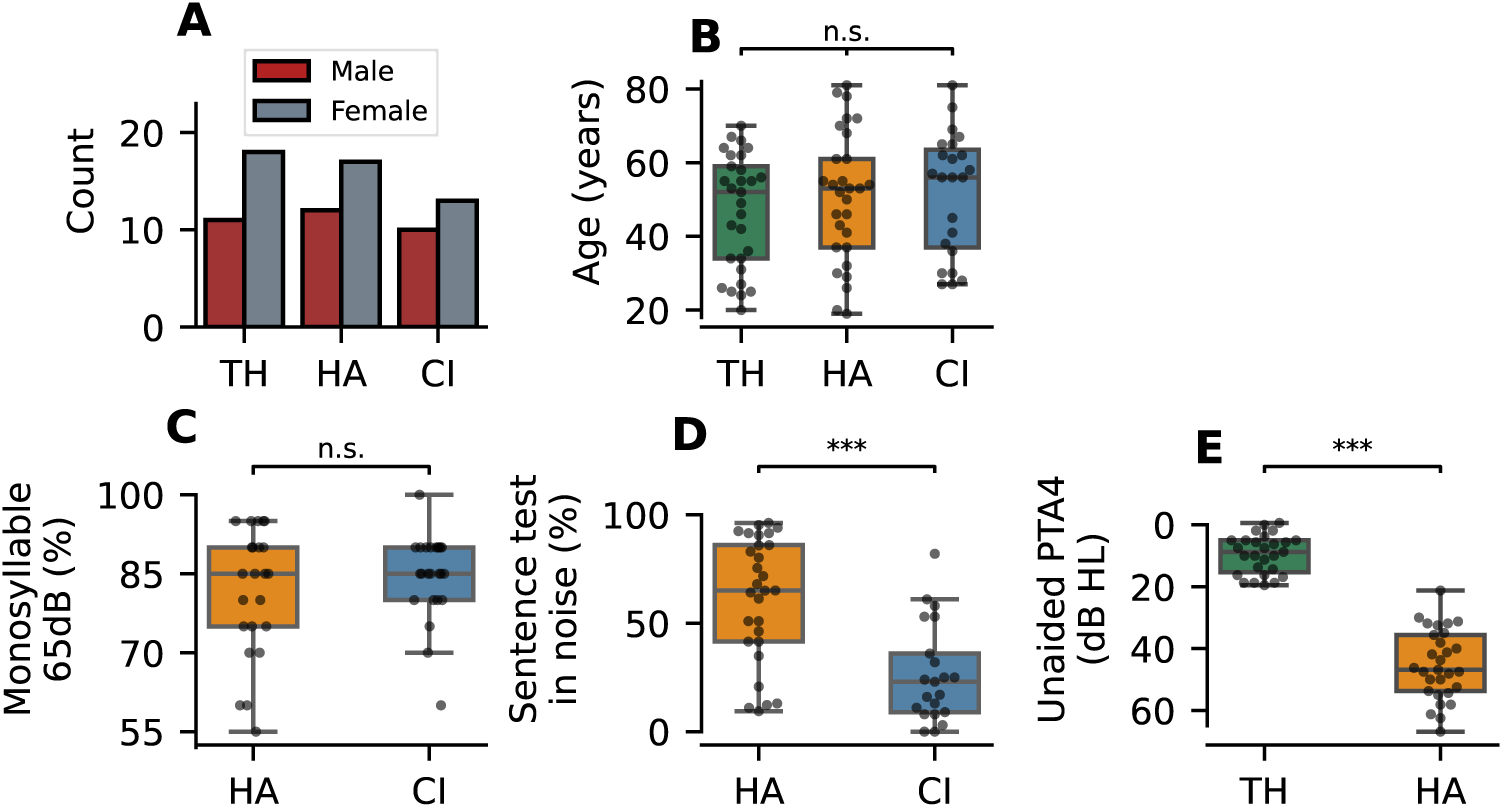
Demographics and audiometric profiles of the three cohorts: 29 typical hearing (TH) participants, 29 hearing aid users (HA), and 24 cochlear implant users (CI). (**A**) All groups show a slight female predominance, with no significant differences in sex distribution across groups. (**B**) Age distributions are broad across cohorts; age-matched recruitment resulted in no significant group differences. (**C**) In the Freiburg monosyllabic word test in quiet (65 dB), both the HA (aided) and CI group achieved a median score of 85%, with no significant difference. (**D**) In the Hochmair-Schulz-Moser (HSM) sentence-in-noise test (65 dB signal, 60 dB noise), the HA group (aided) performed significantly better than the CI group, giving a first indication of better speech-in-noise understanding of the HA group. (**E**) Pure-tone average audiometry results: the TH group had a median PTA4 of 8.75 dB, with all individual values below 25 dB, fulfilling the inclusion criterion. The HA group (unaided) had a median of 46.9 dB, reflecting significantly elevated thresholds. Significance notation: ***:*p <* 0.001, **: *p <* 0.01, *: *p <* 0.05, n.s.: *p >* 0.05.

The Freiburg monosyllabic word test and the HSM test were administered only to the HA and CI groups, as TH individuals would consistently achieve ceiling performance. We confirmed this by testing three TH participants, who all scored 100% in both tests. Panel **C** displays results from the Freiburg monosyllabic word test in quiet, measured at 65 dB with the better ear. The HA and the CI group performed similarly, evident by both their identical median scores of 85% and a two-sided Mann–Whitney U test (MWU) (*U* = 252.5, *p* = 0.796, *r*_rb_ = 0.05). The low rank-biserial correlation (*r*_rb_) indicates that there is almost no statistical evidence in favor of any of the two groups.

Test scores from the HSM sentence-in-noise test (65 dB signal, 60 dB noise) are depicted in panel **D**. In this condition, the HA group (median = 65.1%) performed significantly better than the CI group (median = 23.0%), as confirmed by a two-sided MWU test (*U* = 502.0, *p <* 0.001, *r*_rb_ = 0.64). The high rank-biserial correlation of 0.64 indicates that the vast majority of evidence favors the HA group. This finding provides the first indication within the present study that differences in perception between HA and CI users emerge specifically in the presence of noise.

Finally, unaided pure-tone average thresholds were collected for the TH and HA groups (PTA4: thresholds collected for 0.5, 1, 2, and 4 kHz and then averaged). Panel **E** shows PTA4 values averaged across both ears. As expected the HA group exhibited elevated unaided thresholds (median = 46.9 dB) compared to the TH group. A two-sided MWU test confirmed a significant difference between the HA (*n* = 29) and the TH group (*n* = 29) (*U* = 0.0, *p <* 0.001, *r*_rb_ = 1.0). The maximal rank-biserial correlation of 1 indicates that the entirety of PTA4 scores was elevated in the HA group compared to the TH group.

### 2.2 Stimuli and experimental procedure

For the present study, we employed a selective auditory attention experiment. The stimuli consisted of two German audiobooks: “Elbenwald – Blatt von Tüftler” by J.R.R. Tolkien, narrated by Gert Heidenreich (male) (Tolkien, 2009) and “Eine Frau erlebt die Polarnacht” by Christine Ritter, narrated by Vera Teltz (female) (Ritter, 2020). To ensure comparable speech rates, the speed of the male story was adjusted to 90% of its original speed.

As we are investigating the impact of target speaker identity on behavioral and neural measures, we further characterized our stimuli. To this end, we computed the fundamental frequency (F0) using the pYIN algorithm (*f_min_* = 65 Hz, *f_max_*= 2093 Hz) and the spectral centroids of both speakers (FFT window size = 40 ms, hop length = 20 ms). For both metrics, we trimmed silent parts of the audio and used the librosa package (v0.11.0). Moreover, we estimated the vocal tract length (VTL) of both speakers using a recently developed web application (Anikin et al., 2024). The results, summarized in Table 2, reflect typical differences between male and female voices. The female narrator has a higher fundamental frequency, higher spectral centroids and a shorter vocal tract length.

**Table 2:** Acoustic features for male and female speakers. Values for F0 and Spectral Centroid are reported as mean *±* standard deviation.

| Speaker | F0 (Hz) | Spectral Centroid (kHz) | VTL (cm) |
| --- | --- | --- | --- |
| Male | 110.9 ± 32.5 | 3.46 ± 1.90 | 16.7 |
| Female | 164.5 ± 46.3 | 4.27 ± 2.87 | 14.2 |

Prior to stimulus presentation, both excerpts were normalized to an integrated Loudness Units relative to Full Scale (LUFS) level of *−*24. LUFS is a perceptual loudness metric defined by the ITU-R BS.1770 standard, designed to quantify audio loudness as perceived by the human ear (Ronan et al., 2016).

The stimuli were presented in a free-field environment over two loudspeakers, separated by 60° in azimuth, as schematically depicted in Fig. 2. Both the CI users and the HA users had their hearing devices set to the configuration they typically use in everyday situations. Each participant completed 20 trials, divided into two experimental conditions:

- 8 **single-speaker trials**, in which one audiobook was presented from one loudspeaker alone (either male or female voice), and
- 12 **competing speaker trials**, in which both audiobooks were played simultaneously from separate loudspeakers.

**Fig. 2.**
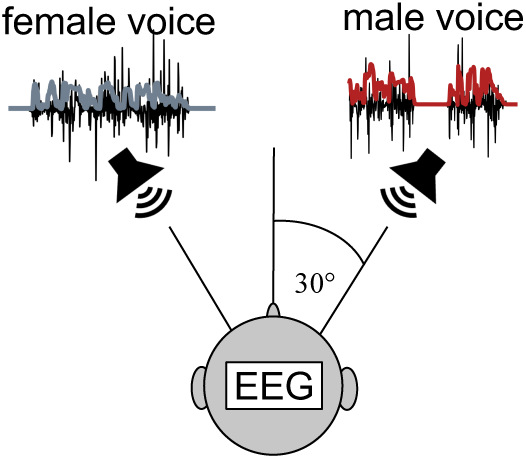
Selective attention paradigm. Two audiobooks were presented over two loudspeakers separated by 60^◦^. Participants were instructed to focus on one of the two audiobooks. The target audiobook was visually communicated over a screen placed centrally before them. Simultaneously, EEG was measured at 1 kHz. Figure adapted from Jehn et al. (2025).

Each trial lasted approximately two minutes. In the competing speaker trials, a visual cue in the form of a symbolic image indicated which audiobook participants should be attending. To facilitate selective attention, the target speaker began 10 s before the distractor. The attended speaker alternated every second trial, and the side of presentation alternated each trial, ensuring counterbalancing. The starting audiobook was pseudorandomized for each participant. The visual instruction was given on a screen in front of the participant, and the participants were instructed to keep their heads aligned with the screen that was placed centrally between the speakers. Adherence was monitored throughout the study.

### 2.3 Behavioral measures

#### Comprehension score

After each trial, participants answered two three-alternative forced-choice (3AFC) questions displayed on the screen before them. These questions were designed to ensure that participants remained attentive to the audiobook content and to assess their level of comprehension under the different listening conditions. The comprehension score was calculated as the ratio of correctly answered questions (*n*_corr_) to the total number of questions (*n*_total_), i.e., *n*_corr_*/n*_total_.

#### Listening effort

Subjective listening effort was evaluated using the ACALES scale, which ranges from 1 (no effort) to 13 (extreme effort) (Krueger et al., 2017). To maintain a manageable experiment duration for participants, effort ratings were collected only once per condition (single vs. competing speaker scenario) and attended speaker (male or female). Participants were shown a visual scale from 1 to 13, including descriptive labels for each level, and verbally reported their perceived listening effort to the experimenter.

### 2.4 EEG recordings

EEG data were recorded using an actiCHamp system (BrainProducts GmbH, Germany) with a total of 32 electrodes. Of these, one was positioned below the eye to facilitate the detection and removal of putative ocular artifacts, resulting in 31 electrodes being used to capture neural activity. EEG was measured at a sampling rate of 1 kHz, and the signal was online low-pass filtered with a cut-off frequency of 280 Hz. Electrode impedances were assessed both before the recording and during the break between the single and competing speaker scenario with a 1 *V pp* excitation signal at a frequency of 30 Hz. Impedances were kept below 20 kΩ for the entire recording session.

In CI users, up to two electrode positions were occasionally located directly between the implant and the CI coil, impeding reliable recordings at those sites. In those cases, the affected electrodes were removed from the cap. Across both hemispheres, an average of 1.7 electrodes per CI user were removed.

To synchronize the audio stimulus with the EEG recordings, the presented audio signal was split and simultaneously recorded as two auxiliary channels via the EEG system. This was achieved using two Stim-Trak adapters (BrainProducts GmbH, Germany) connected through an audio splitter. An offline correlation analysis was conducted between the recorded audio signal and time-shifted versions of the clean stimulus waveforms. The delay corresponding to the maximal Pearson correlation coefficient was selected to temporally align each stimulus with the EEG data. Onset triggers were also transmitted at the start of each stimulus presentation and served as a validation reference for the computed delays.

### 2.5 Pre-processing

#### Audio stimuli

Speech envelopes were extracted by applying the Hilbert transform to the original audio signals (48 kHz) of both speech streams, followed by taking the absolute value of the resulting analytic signals. In accordance with Nogueira and Dolhopiatenko (2022), the auditory periphery was not modeled during envelope extraction due to the limited spectral resolution inherent to CIs. To maintain consistency in preprocessing, the same approach was applied to both the TH and HA groups. The resulting envelopes were low-pass filtered below 8 Hz (one-pass, zero-phase, non-causal type 1 FIR filter, *−*6 dB cutoff frequency: 9.0 Hz, order 79 200, Hamming window with 0.0194 passband ripple and 53 dB stopband attenuation) and resampled to 128 Hz. Then the envelopes were high-pass filtered above 1 Hz (one-pass, zero-phase, non-causal type 1 FIR filter,*−*6 dB cutoff frequency: 0.5 Hz, order 422, Hamming window with 0.0194 passband ripple and 53 dB stopband attenuation).

#### EEG

Missing EEG channels in the CI cohort due to the implant (see Section 2.4) were estimated from neighboring electrodes using spline interpolation. EEG recordings were first low-pass filtered offline with an upper passband edge of 8 Hz (one-pass, zero-phase, non-causal type 1 FIR filter, *−*6 dB cutoff frequency: 9.0 Hz, order 1650, Hamming window with 0.0194 passband ripple and 53 dB stopband attenuation) and then downsampled to 128 Hz. Subsequently, a high-pass filter with a lower passband of 1 Hz (one-pass, zero-phase, non-causal type 1 FIR filter, *−*6 dB cutoff frequency: 0.5 Hz, order 422, Hamming window with 0.0194 passband ripple and 53 dB stopband attenuation), was applied to mitigate slow drifts. Both filters were designed using the default settings from the MNE Python library (version 1.5.1) (Gramfort et al., 2013), which implements the suggestions for EEG filtering made by Widmann et al. Widmann et al. (2015). After filtering, all EEG channels were standardized to achieve zero mean and unit variance per trial. The reference was set to the average of all recorded EEG channels.

#### Backward modeling

CI-related artifacts were not explicitly removed in the backward model, as the model can learn to downweight affected channels during training and is thus generally robust to spatially localized noise. In previous work on the same CI dataset, it has been shown that additional artifact rejection did not improve decoding accuracy for the linear backward model Jehn et al. (2025). Moreover, manual inter-rater variability in artifact labeling leads to a lack of reproducibility (Delorme, 2023). Therefore, no additional artifact rejection was applied in the linear backward model.

#### Forward modeling

Unlike backward models, which are evaluated on reconstruction scores and decoding accuracy, our forward model analysis involves visualizing model weights and topographies. Because these visualizations are sensitive to artifacts, we removed physiological and CI artifacts to prevent them from masking the neural signal and confounding the interpretation. Especially artifacts introduced by the CI are problematic, as their magnitudes can easily exceed the neural signal (Gilley et al., 2006; Viola et al., 2012).

We removed physiological artifacts in datasets from all cohorts, using the MNE implementation of the automatic ICLabel framework (Pion-Tonachini et al., 2019). To this end, the EEG data sampled at 1 kHz was band-pass filtered between 1 and 100 Hz (one-pass, zero-phase, non-causal type 1 FIR filter, *−*6 dB lower cutoff frequency: 0.5 Hz, *−*6 dB upper cutoff frequency: 112.5 Hz, order 3300, Hamming window with 0.0194 passband ripple and 53 dB stopband attenuation). This configuration aligns with the frequency range recommended by MNE for optimal classification performance. ICA was then applied using the infomax algorithm, separating 30 components per trial. Components classified by the ICLabel algorithm as “eye”, “muscle”, “heart”, or “line noise” with a probability greater than 50% were subsequently removed. Finally, the cleaned data were low-pass filtered below 8 Hz and downsampled to 128 Hz, as described in the previous EEG paragraph. The number of components rejected as physiological artifacts per trial was 6.1 *±* 4.3 for the TH cohort, 4.1 *±* 1.8 for the HA cohort, and 11.1 *±* 3.6 for the CI cohort.

To suppress the presence of CI artifacts and allow interpretable data visualization, we employed an automated approach that is based on Independent Component Analysis (ICA). Specifically, we followed the methodology proposed in (Jehn et al., 2025), where the time series of each IC of the EEG is cross-correlated with the audio signal. First, an ICA with 30 components was computed using the InfoMax algorithm (Bell and Sejnowski, 1995). The similarity of an EEG component to the audio stimuli was quantified by the signal-to-noise-ratio (SNR) of the peak in the cross-correlation function around a delay of 0 s (*±*5 ms) in dB. As cortical responses emerge only later, high SNR values indicate signals that emerged from the CI stimulation. Components with an SNR above 15 dB were rejected as CI artifacts. On average, 11.1 *±* 4.9 components were discarded in this manner.

### 2.6 Linear backward model

Linear backward models, also known as linear decoders, are trained to reconstruct the presented speech envelope *y*(*t_i_*) at time *t_i_* from brain signals, as acquired from a *J*-channel EEG recording 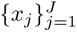. When a participant is focusing on a specific voice in a multi-talker situation, the speech envelope reconstructed by the backward model 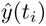 is more similar to the envelope of the attended signal than to other present signals (Etard et al., 2019; O’Sullivan et al., 2015).

#### 2.6.1 Metrics

The similarity can be quantified by calculating the Pearson correlation coefficient between the reconstructed speech envelope *y*^ and the envelope of the attended audio *y_att_* as

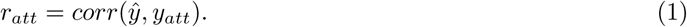

In the same way, the Pearson correlation coefficient between the reconstructed speech envelope *y*^ and the envelope of the ignored audio *y_ign_* is obtained as

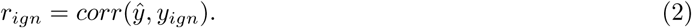

We refer to these correlation coefficients as reconstruction scores; the larger one is taken as an indication that the corresponding voice is attended. If *r_att_* is consistently greater than *r_ign_*, auditory attention can be successfully decoded. To evaluate this, decoding accuracy is computed by calculating *r_att_* and *r_ign_* for discrete segments of data. From the number of correctly classified segments *n_corr_*, where *r_att_ > r_ign_* holds, and the total number of considered segments *n_total_*, the decoding accuracy is computed as

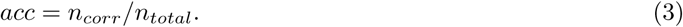

#### 2.6.2 Model definition

In the present study, we used a linear ridge regression model where the reconstructed envelope at time *t_i_* is computed as

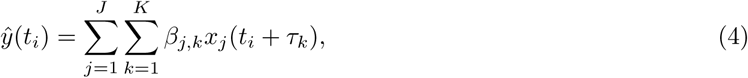

with learnable model parameters *β_j,k_*. For the linear backward models, we considered *K* = 128 time lags

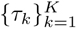 ranging from *−*500 ms to 500 ms. The parameters of the linear model were optimized by minimizing the L2-regularized sum of squared errors *E*(*β_j,k_*) between the ground truth envelope *y*(*t_i_*) and the model estimate 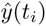. The loss function is thus expressed as

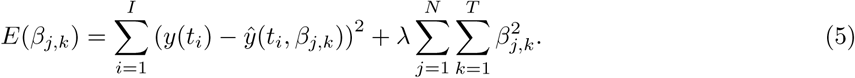

The regularization term 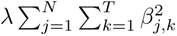, penalizes large parameter values in *β_j,k_*, and therefore counteracts overfitting. The hyperparameter *λ* controls the impact of the regularizer and was optimized on a held-out validation set based on maximal reconstruction scores. Tested hyperparameters *λ* ranged logarithmically from 10^−7^ to 10^7^.

We trained individual models for each subject using twelve-fold cross-validation. In each fold, one competing-speaker trial was designated as the test set, while another trial—pseudo-randomly selected—served as the validation set for tuning the regularization parameter. The remaining 18 trials were used to fit the regression model.

#### 2.6.3 Explanatory modeling of neural metrics

To investigate whether age, level of hearing loss, or behavioral responses influenced cortical envelope tracking across participant groups, we fitted linear mixed-effects models (LMEMs) separately for each group. Only data from competing speaker trials were included in the analysis, as these most closely reflect real-world cocktail party listening scenarios, and both attended and ignored reconstruction scores were available.

Cortical tracking was quantified using three response variables (denoted *y* in our LMEMs) (1) the reconstruction score of the attended envelope (*r_att_*), (2) the reconstruction score of the ignored envelope (*r_ign_*), and (3) the decoding accuracy from 30-second decision windows. Data were analyzed at the trial level to capture trial-by-trial variability in comprehension and its relationship with the response variables.

Fixed effects included listening effort (*le*) and age, modeled across all cohorts. To account for repeated measurements within individuals, a random intercept was included for each subject. Subject-level predictors, such as age, were constant across trials and replicated accordingly. Due to significant ceiling effects, the comprehension score (*cs*) was included as a predictor exclusively for the CI cohort, as this measure lacked sufficient variance for meaningful analysis in the TH and HA groups. Each predictor was standardized (z-scored) prior to model fitting to facilitate interpretation of model coefficients. To assess the assumption of normality of residuals, Q–Q plots were examined for each fitted model.

Within the TH group, we additionally examined the effect of hearing loss, quantified as unaided pure-tone average (*pta4*; see Fig. 1). Given the interdependence between age and hearing loss, an interaction term between these two variables was included to capture their joint influence. The resulting linear mixed-effects model was specified as:

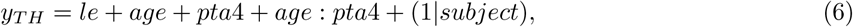

where *y_T H_* denotes the response variable within the TH group.

For the HA group, we extended the model to include additional measures of speech perception: the Freiburg monosyllabic word test in quiet at 65 dB (*word*) and the Hochmair-Schulz-Moser sentence-in-noise test (*hsm*). These predictors were incorporated alongside behavioral and demographic variables in a linear mixed-effects model defined as:

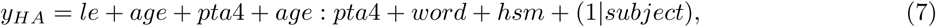

where *y_HA_* denotes the response variable within the HA group, and a random intercept was included for each subject to account for repeated measures. For five HA participants, the Freiburg monosyllabic word test was performed in a non-standardized way These data were treated as missing values.

For the CI group, unaided PTA4 values could not be obtained and were therefore not included in the analysis. As a result, hearing loss was not considered in the linear mixed-effects model, which was specified as:

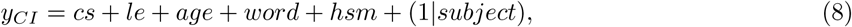

where *y_CI_* denotes the response variable within the CI group, and a random intercept was included for each subject to account for repeated measures.

P-values for each predictor were corrected for multiple comparisons using FDR correction, based on the number of models in which that predictor appeared. For example, comprehension score (cs) was included in 9 separate models, while hearing loss (PTA4) was included in 6 models; FDR correction was applied accordingly for each predictor.

### 2.7 Speaker bias

In a previous study on the identical CI dataset, a preference among CI users, as quantified by listening effort and comprehension score, for the female speaker was observed (Jehn et al., 2025). This behavioral preference was mirrored in the neural data, where the reconstruction scores from the backward model showed a systematic bias toward the female speaker. In the present study, we recorded data from both HA and TH individuals using the same experimental setup. Our objective was to determine whether the observed bias was an inherent feature of the setup itself or a phenomenon specific to the CI group.

First, we separated listening effort ratings and comprehension scores for each of the two voices and examined whether behavioral differences emerged between the two speakers. Second, we computed reconstruction scores for the male speaker (*r*_male_) and the female speaker (*r*_female_) on segments of 60 s in duration, yielding pairs (*r*_male_*, r*_female_). To assess a potential bias in the reconstruction scores toward either the male or female speaker, we computed the percentage of segments where *r_female_ > r_male_*. This value should approach 50% for a balanced decoder.

### 2.8 Linear forward model

A linear forward model, or encoder, is trained to predict the neural response from the presented stimulus. Unlike backward models, forward models enable neurophysiological interpretation of the weights, known as the temporal response function (TRF) (Crosse et al., 2021). It is well established that auditory processing in humans is hierarchically organized, starting at the auditory nerve and continuing to primary and non-primary auditory cortex (Davis and Johnsrude, 2003; O’Sullivan et al., 2019). Early TRF components in multi-talker situations (30 ms - 80 ms), localized in Heschl’s gyrus (HG), mainly capture acoustic processing of the stimulus and are not substantially modulated by attention (O’Sullivan et al., 2019; Puvvada and Simon, 2017). In contrast, later responses (*>* 85 ms) in the superior temporal gyrus (STG), are heavily modulated by attention and mainly carry information of the attended talker (Golumbic et al., 2013; O’Sullivan et al., 2019). By analyzing the TRFs of hearing assistive technology users, we can draw conclusions about the selectivity of cortical processing at different stages of their auditory pathway.

#### Model definition

We computed forward models that aim to infer the *J*-channel EEG recording 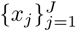 based on the presented speech envelope *y*(*t_i_*) at time *t_i_*. A single EEG channel is predicted as

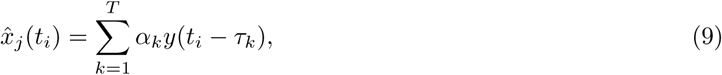

for 141 equispaced time lags *τ_k_*ranging from -300 to 800 ms (i.e. 7.8 ms between lags) and learnable model parameters *α_k_*. The parameters were estimated using ridge regression with the error term

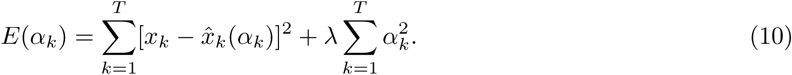

#### Training

Regularization parameters *λ*, logarithmically spaced from 10^−5^ to 10^5^, were evaluated at the population level across all cohorts. Using fivefold cross-validation, *λ* = 1 consistently yielded the highest encoding scores, defined as the Pearson correlation between the predicted neural signals 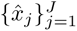 and the recorded neural signals 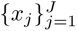, averaged across all EEG channels. Adopting a common regularization parameter across all groups enables a fair comparison of TRFs between cohorts. TRFs were computed at the trial level, which avoids introducing noise caused by the concatenation of trials. One model was computed for each competing speaker trial and the model coefficients were averaged for each subject. In each trial, 20% of the data were left out of training to compute the encoding score to quantify the predictive power of the model. To study the effect of attention in the three groups, we computed TRFs for the attended and ignored speech envelopes and compared their encoding scores and model coefficients.

#### Null model

To validate the integrity of the computed TRFs, we generated null models by training 500 forward models on circularly time-shifted versions of the presented stimuli. The time shifts were uniformly spaced between one-third and two-thirds of the trial length, ensuring that the stimulus-EEG phase relationship was effectively disrupted while preserving the autocorrelated structure of both signals. Each null model yielded a null TRF with a misaligned stimulus-response phase. Their encoding scores form the null distribution that is expected to be centered around 0. We then compared the encoding scores of the attended and ignored TRFs against this null distribution, using statistical tests and correction for multiple comparisons as described in section 2.9. If the encoding scores of the attended and ignored TRFs significantly exceed those of the null distribution, this provides evidence for their validity.

#### Coefficient post-processing

Although the EEG data were average-referenced, the grand average of the CI group displayed a pronounced low-frequency drift that superimposed local variations in the TRF. To address this and facilitate accurate interpretation, we subtracted the mean across channels at each time lag *τ_k_*. For consistency, the same post-processing procedure was applied to all groups. We computed the grand average TRF by averaging the coefficients across all subjects within a group. We then calculated the magnitude of the grand average TRF, obtained by taking the absolute value of each channel’s response, followed by averaging across channels.

#### Significant lags

To identify significant time lags, we adopted the method by (Kegler et al., 2022). To this end, we compared the magnitudes of the grand average magnitude TRF for the attended envelope against a null distribution. This null distribution consisted of 500 grand average TRFs generated as described in the “Null model” paragraph. For each time lag, we calculated an empirical p-value by finding the proportion of null TRFs whose magnitude exceeded that of the attended TRF. These p-values were then corrected for multiple comparisons using the Bonferroni method. A time lag was considered statistically significant if its corrected p-value was less than 0.05.

#### Statistical analysis of coefficients

To compare attended and ignored responses, we first computed a TRF for both conditions (attended, ignored) per subject. For each subject, TRFs for the attended and ignored TRFs were derived by averaging models computed separately on each of the twelve competing speaker trials. We then calculated the magnitude TRFs of each subject by taking the absolute value of each channel, followed by averaging across channels. To reduce the influence of local noise, we averaged TRF magnitude values within a 7 ms window centered around the global attended peak. This procedure yielded distributions of attended and ignored TRF magnitudes, which were then statistically compared as described in Section 2.9.

### 2.9 General statistical procedures

#### Comparative statistical testing

Statistical comparisons were performed at the subject level, with trial-level values averaged per participant to avoid within-subject dependence and ensure consistency in the treatment of subject-level predictors such as age, listening effort, and hearing thresholds. This subject-level approach inherently accounts for within-subject variance, thus precluding the need for random intercepts, which are a necessary component of our trial-level neural-behavior analysis. Normality and homogeneity of variance were assessed using the Shapiro–Wilk test and Levene’s test, respectively.

For between-group comparisons, a one-way ANOVA followed by post hoc unpaired t-tests was applied when assumptions were met. Otherwise, the Kruskal–Wallis test was used, followed by post-hoc Mann–Whitney U tests.

For within-group comparisons (e.g., comparisons of encoding scores from different models on the same cohort), repeated-measures ANOVA followed by post hoc paired t-tests was applied when assumptions were satisfied. Otherwise, the Friedman test was used as a non-parametric alternative, followed by Wilcoxon signed-rank tests for pairwise comparisons.

Post hoc p-values were corrected for multiple comparisons using the Benjamini–Hochberg False Discovery Rate (FDR) procedure.

#### Chance level estimation for comprehension scores

We used two-sided binomial tests to determine whether comprehension scores exceeded chance-level performance (33%). Tests were performed at the trial level by comparing the number of correctly answered questions to the expected chance level. We then identified the threshold for above-chance performance based on the 95% confidence level and visualized it in the corresponding plots.

#### Significance level and thresholds of reporting

For all statistical tests, we used a significance threshold of *α* = 0.05. In figures, statistical significance is indicated as follows: ^∗∗∗^ for *p <* 0.001, ^∗∗^ for *p <* 0.01, ^∗^ for *p <* 0.05, and n.s. for non-significant results.

### 2.10 Software

All analyses were conducted in Python 3.9.7. Comparative statistical testing and multiple comparison corrections were performed using the pingouin package (v0.5.5), while binomial tests were computed with scipy (v1.12.0). Linear mixed-effects models were fitted using statsmodels (v0.14.2). Linear forward and backward modeling was implemented with the sPyEEG (v0.0.1) package (Guilleminot et al., 2021). EEG preprocessing and topographical visualizations were carried out using MNE (v1.8.0) (Gramfort et al., 2013). Figures were generated with matplotlib (v3.7.0) and seaborn (v0.13.2). The source code for data analysis is available under https://github.com/Constantin-Jehn/aad-neuroimage.git.

## 3 Results

### 3.1 Behavior

In Fig. 3, we present the participants’ listening effort ratings (panels **A** and **B**), where higher values correspond to greater perceived effort, and their comprehension scores (panels **C** and **D**), calculated as the percentage of correctly answered questions. Each data point in the panels represents a participant’s average listening effort rating or comprehension score. In all panels, at least one cohort exhibited a non-normal distribution, necessitating the use of non-parametric tests for the analysis of behavioral scores.

**Fig. 3.**
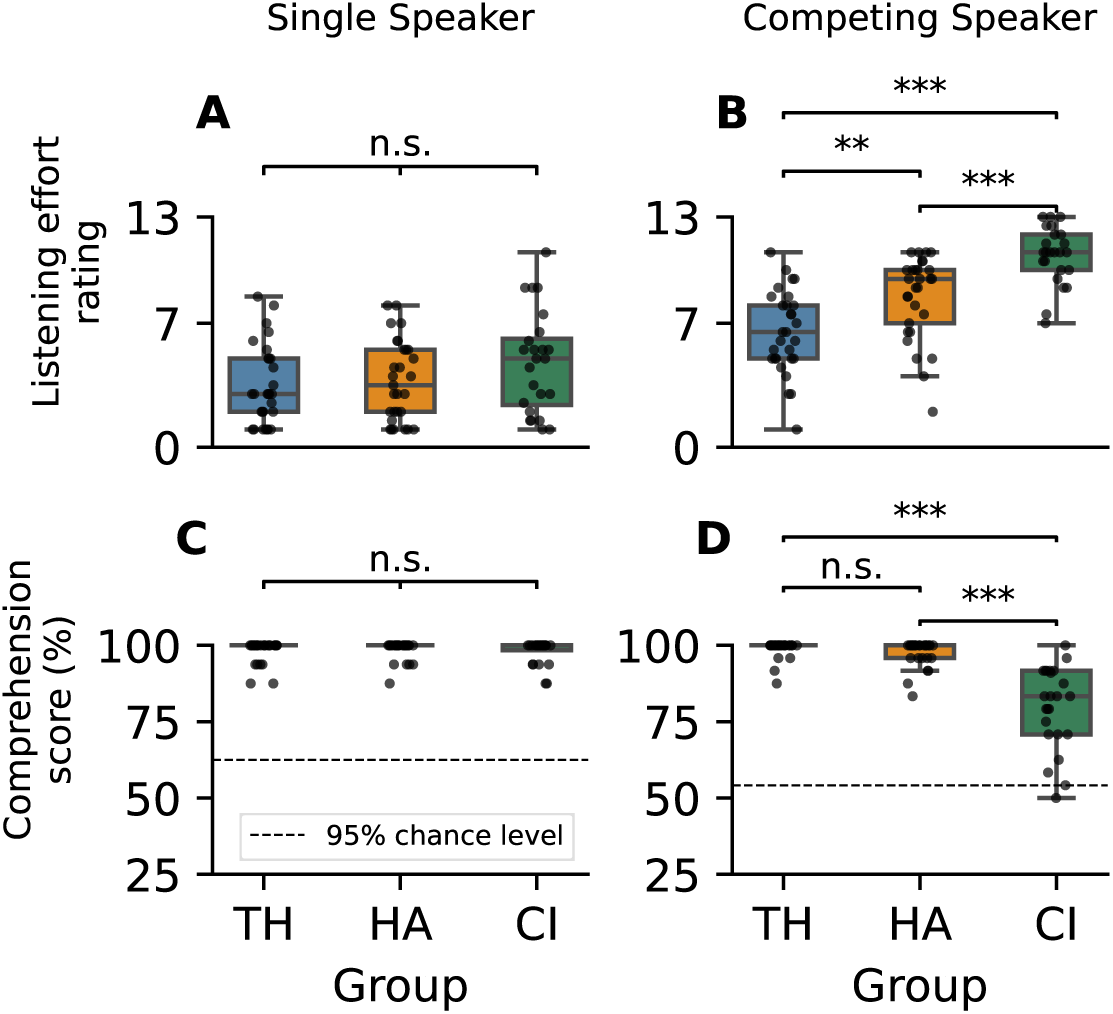
Listening effort and comprehension score under single-speaker and competing-speaker conditions. (**A**) The median listening effort was rated from 3 (“very little”) to 5 (“little”) in the single speaker paradigm, with several upper outliers in the CI group. No significant group difference emerged. (B) Listening efforts were increased across groups under the competing speaker condition. The median ratings ranged from 6.5 (“considerable effort”) in the TH group to 11.0 (“very much effort”) in the CI group. The stepwise increase in listening effort from the TH to HA to CI group was statistically significant. (C) Under the single speaker condition, the comprehension scores of all groups were at ceiling, without significant group differences. (**D**) While most participants in the TH and HA cohorts maintained perfect scores (100%) in the competing-speaker condition, the median comprehension score in the CI group dropped to 83%. The reduction in scores for the CI group compared to the TH and HA groups was significant, whereas no significant difference was found between the TH and HA cohorts. Significance notation: ***:*p <* 0.001, **: *p <* 0.01, *: *p <* 0.05, n.s.: *p >* 0.05.

#### 3.1.1 Listening effort

In the single-speaker paradigm, the median listening effort ratings of the TH group (median = 3.0) and the HA group (median = 3.5) correspond to “very little effort”, whereas the CI group’s median rating of 5.0 reflects “little effort” (Panel A). Notably, several individuals from the CI group reported high listening effort already in the simpler condition. Despite the elevated median in the CI group, a Kruskal–Wallis test indicated no significant group differences in listening effort in the single-speaker scenario (*H*(2) = 3.31, *p* = 0.190). The small effect size (*η*^2^ = 0.016) confirmed that group membership had little practical impact on listening effort in this condition. As expected, listening effort increased substantially in the competing-speaker paradigm (Panel B). Median ratings rose to “moderate effort” in the TH group (median = 6.5), “considerable effort” in the HA group (median = 9.5), and “very much effort” in the CI group (median = 11.0). A Kruskal–Wallis test confirmed significant differences between groups (*H*(2) = 36.27, *p <* 0.001, *η*^2^ = 0.43). The large effect size indicated substantial differences in perceived effort between the cohorts. Post-hoc two-sided Mann–Whitney U (MWU) tests revealed significantly greater listening effort in the HA group compared to the TH group (*U* = 624.0, *p* = 0.002, *r*_rb_ = 0.48), and higher effort in the CI group relative to the HA group (*U* = 570.0, *p <* 0.001, *r*_rb_ = 0.64). Unsurprisingly, the difference between the CI and TH group was also significant (*U* = 652.5, *p <* 0.001, *r*_rb_ = 0.88). All effect sizes (rank-biserial correlation) of the inter-group comparisons were above 0.45 and thus indicated substantial differences between all groups. In this case, p-values were corrected for three post-hoc comparisons using the FDR method.

Note that the listening effort in the competing speaker paradigm was comparable between post-lingually implanted CI users (n=20) (median = 10.85, range = 7.0 *−* 13.0) and the pre- and peri-lingually implanted CI users (n=4) (median = 11.25, range = 10.5 *−* 12.0). A one-way ANOVA confirmed that there was no significant influence of the time of implantation (*F* = 0.164, *p* = 0.84).

#### 3.1.2 Comprehension score

The lower row in Fig. 3 displays the measured comprehension scores. Panel **C** shows that all groups performed at ceiling in the *single-speaker* paradigm, with a median comprehension score of 100% across all cohorts. A Kruskal–Wallis test confirmed the absence of significant group differences (*H*(2) = 0.53, *p* = 0.764, *η*^2^ = 0.02). In the *competing-speaker* scenario, both the TH and HA groups continued to perform at ceiling, with median comprehension scores of 100% (Panel **D**). Only four participants in the TH group and nine in the HA group failed to achieve a perfect score. In contrast, the CI group showed markedly reduced performance, with a median score of 83% and with four participants scoring close to the 95% confidence interval (corresponding to 54% correct answers). A Kruskal-Wallis test demonstrated a significant group difference in comprehension in the competing-speaker condition (*H*(2) = 49.8, *p <* 0.001, *η*^2^ = 0.61). The effect size was very large, indicating that over 60% of the variance in comprehension scores can be attributed to group membership.

No significant difference was observed between the TH and HA cohort (two-sided MWU: *U* = 348.5, *p* = 0.127), with a small effect size (*r*_rb_ = *−*0.17). However, post-hoc two-sided MWU tests indicated that scores were significantly deteriorated in the CI group, with very large effect sizes observed for both comparisons (TH vs. CI: *U* = 28.5, *p <* 0.001, *r*_rb_ = *−*0.92 and HA vs. CI: *U* = 46.0, *p <* 0.001, *r*_rb_ = *−*0.87). Note that the post-hoc tests were FDR-corrected for three comparisons and that the 95% chance level in the single speaker is higher because fewer trials were conducted in this condition.

The comprehension scores in the competing speaker paradigm were slightly increased in the post-lingually implanted CI users (n=20) (median = 0.833, range = 0.46 *−* 1.0) compared to the pre- and peri-lingually implanted CI users (n=4) (median = 0.75, range = 0.58 *−* 0.96), but a one-way ANOVA found no significant difference across groups (*F* = 0.181, *p* = 0.85).

### 3.2 Neural decoding

Our primary objective was to study cortical speech tracking in individuals using hearing assistive technologies, i.e., HAs or CIs. To this end, we first trained linear backward models for the three cohorts and compared model results in Fig. 4. For maximal objectivity of the analysis, we applied only filtering between 1 and 8 Hz to the EEG data without additional pre-processing.

**Fig. 4.**
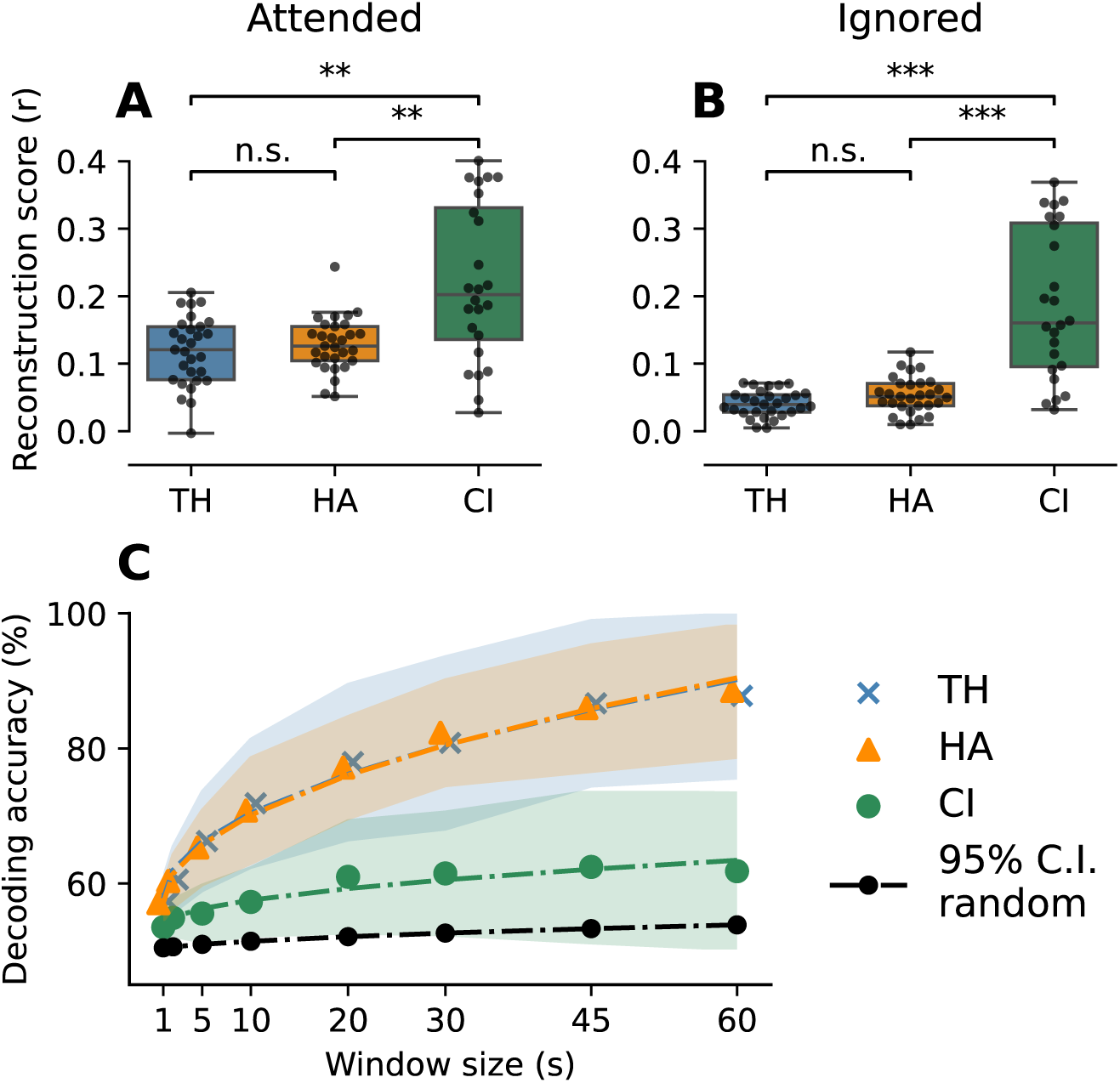
Results from the linear backward model. (**A**) Attended reconstruction scores were similar in the TH (median = 0.121) and HA (median = 0.127) group. The CI group exhibited significantly increased reconstruction scores (median = 0.194), which can be mainly attributed to the presence of electrical artifacts. (**B**) Ignored reconstruction scores were substantially lower in the TH (median = 0.038) and the HA (median = 0.051) group compared to their respective attended scores. The CI group (median = 0.157) showed decreased reconstruction scores for the ignored speaker as well. Between-group differences were only significant between the CI and TH/HA group. (**C**) All groups showed declining decoding accuracies with shorter window sizes but remained above the chance level. While the TH and HA groups performed comparably, reaching maximal decoding accuracies at 60 s of 87.8% and 88.5% respectively, the CI group performed consistently worse and scored a maximal accuracy of 63.1%. Significance notation: ***:*p <* 0.001, **: *p <* 0.01, *: *p <* 0.05, n.s.: *p >* 0.05.

#### 3.2.1 Reconstruction scores

Backward models were trained to reconstruct the attended envelope *y* from acquired EEG data. Reconstruction scores were computed as the Pearson correlation coefficient *r_att_* between the reconstructed envelope *y*^ and the actual envelope of the attended speaker *y_att_*(Panel **A**) as well as the correlation coefficient *r_ign_* between the reconstructed envelope *y*^ and the envelope of the ignored speaker *y_ign_*(Panel **B**). Each data point in Panels A and B represents a participant’s average score determined by twelve-fold cross-validation, with test sets drawn from the competing speaker condition. The reconstruction scores were computed on 60 s segments. As homogeneity of variances neither held for attended nor for ignored reconstruction scores, non-parametric tests were employed. All p-values from post-hoc comparisons were corrected for multiple testing using the FDR method for three comparisons.

In Panel **A**, we observe similar *attended* reconstruction scores in the TH group (median = 0.121) and the HA group (median = 0.126). The distribution of the CI group (median = 0.202) shows a larger variance with values up to 0.40.

A Kruskal-Wallis test confirmed significant differences between groups (*H*(2) = 13.1, *p* = 0.002), with a medium-to-large effect size (*η*^2^ = 0.14). Post-hoc two-sided MWU tests showed no significant difference between the TH (*n* = 29) and HA (*n* = 29) group (*U* = 384.0, *p* = 0.576), with a negligible effect size (*r*_rb_ = 0.09). In contrast, attended reconstruction scores in the CI group (*n* = 24) were significantly increased compared to the TH group (*U* = 164.0, *p* = 0.003, *r*_rb_ = 0.52) and the HA group (*U* = 188.0, *p* = 0.004, *r*_rb_ = 0.48), indicating large effects for both comparisons. The post-hoc comparisons of attended reconstruction scores were FDR-corrected for three comparisons. The substantially elevated reconstruction scores can be attributed to artifacts introduced by the CI. Since the electrical stimulation impulses are highly correlated with the presented stimuli and their envelopes, they lead to an increase in reconstruction scores produced by the backward model.

Panel **B** displays the reconstruction scores for the *ignored* speech. Both the TH group (median = 0.038) and the HA group (median = 0.051) showed substantially reduced reconstruction scores for the ignored condition compared to the attended one. The CI group likewise exhibited significantly lower reconstruction scores in the ignored condition (median = 0.159) compared to the attended condition (median = 0.202; Wilcoxon signed rank test: *n* = 24; *W* = 1.0, *p <* 0.001), reflecting attentional modulation with a very large effect size (*r*_rb_ = 0.993). However, the modulation was less pronounced in the absolute scores and accompanied by much greater within-group variability than in the other two groups.

We also found a significant group difference in the ignored reconstruction scores between groups (Kruskal-Wallis test: *H*(2) = 34.4, *p <* 0.001), with a large effect size (*η*^2^ = 0.41). The difference between the TH (*n* = 29) and HA (*n* = 29) groups approached, but did not reach, statistical significance (two-sided MWU: *U* = 303.0, *p* = 0.069), showing a small-to-medium effect size (*r*_rb_ = 0.28). Post-hoc two-sided MWU tests confirmed that the differences between the TH and CI group (*U* = 53.0, *p <* 0.001, *r*_rb_ = 0.85), as well as between HA and CI users (*U* = 86.0, *p <* 0.001, *r*_rb_ = 0.75) were statistically significant and accompanied by very large effect sizes. The post-hoc comparisons of ignored reconstruction scores were FDR-corrected for three comparisons.

#### 3.2.2 Decoding accuracy

We can use the reconstruction scores to decode auditory attention on a specific segment of the EEG. As we expect *r_att_* to exceed *r_ign_* due to enhanced cortical tracking of attended speech, a segment is classified correctly if *r_att_ > r_ign_*. From the number of correctly classified segments, *n_corr_*, and the total number of segments, *n_total_*, we obtain the decoding accuracy as *n_corr_/n_total_*. As we vary the length of the segments for decoding, we obtain a curve of decoding accuracy as a function of window size, shown in Fig. 4 **C**. The data points in this plot represent the population average in decoding accuracy, with the values for each subject derived from twelve-fold cross-validation. Shaded areas indicate the population mean *±* one within-group standard deviation. As a reference, the 95% chance-level for attention decoding is also plotted: it slightly increases with a larger window size, as fewer segments can then be classified.

All groups exhibited a decline in decoding accuracy for shorter decision windows, following approximately a square-root function. The data points of the TH and HA groups almost overlap for all window sizes, indicating comparable decoding performance. They reach a maximal average decoding accuracy of 87.8% (TH) and 88.5% (HA) for 60 s windows. These findings are consistent with the similar reconstruction scores observed in Panels A and B. The CI group stands out with consistently lower decoding accuracies across all window sizes, reaching a maximal decoding accuracy of only 62.1% for 60 s time windows. The lower decoding accuracy can be attributed to a weaker attentional modulation of reconstruction scores, as observed in Panels A and B. We suspect that lower levels in speech comprehension (see Fig. 3 **D**) contribute to the diminished decoding accuracy in this group. As the time of implantation is a putative factor influencing the neural segregation, we compared the decoding accuracy (30 s decision window) between post-lingually implanted (n=20) and peri/pre-lingually implanted CI users. The accuracies were found to be indistinguishable between these groups (post-lingual: median = 0.61, range = 0.52 *−* 0.79, peri/pre-lingual(n=4): median = 0.62, range = 0.52*−*0.63). A one-way ANOVA confirmed that there was no influence of time of implantation on decoding accuracy (*F* = 0.176, *p* = 0.84).

#### 3.2.3 Explanatory modeling of neural metrics

##### Comprehension score

In the CI group, higher comprehension scores were associated with increased neural tracking of the attended envelope (*β* = 0.098, 95% CI = [*−*0.003, 0.200], p = 0.087) and decreased tracking of the ignored envelope (*β* = *−*0.087, 95% CI = [*−*0.192, 0.018], p = 0.103). While these effects did not reach statistical significance, coefficients and confidence intervals indicate a potential relationship between comprehension and selective neural tracking. Finally, decoding accuracy, which reflects the difference between *r_att_*and *r_ign_*, increased significantly for participants with higher comprehension scores (accuracy: *β* = 0.236, 95% CI = [0.064, 0.407], p = 0.021). Note that this significant correlation is plausible despite the presence of electrical stimulation artifacts. Unlike the absolute reconstruction scores, decoding accuracy, as defined in equation 3, involves solely the contribution from the neural sources, and not potential CI artifacts, since the latter do not vary with attention. This suggests enhanced neural differentiation between attended and ignored speech in participants with better comprehension.

##### Hearing loss

In the TH group, a greater degree of hearing loss was a significant predictor of increased neural tracking of the ignored envelope (*β* = 0.309, 95% CI = [0.103, 0.514], p = 0.019). Furthermore, hearing loss showed a positive association with neural tracking of the attended envelope (*β* = 0.382, 95% CI = [0.037, 0.727], p = 0.089). Although this latter result did not reach significance, the confidence interval being entirely in the positive domain, suggests a likely positive effect. Decoding accuracy, however, was not significantly predicted by hearing loss (*β* = 0.200, 95% CI = [*−*0.024, 0.421], p = 0.161), which is plausible as both attended and ignored reconstruction scores were positively impacted by greater hearing loss.

In the HA group, unaided PTA4 values were not a significant predictor for any of the outcome measures (all p*>*0.2), which is plausible given that hearing loss was compensated for by hearing aids throughout the experiment.

##### Null result

Interestingly, neither listening effort nor age significantly affected any of the response variables in any of the three groups. Also, the interactions between hearing loss and age were not significant in both the TH and HA group for all response variables, suggesting that the effect of hearing loss on cortical tracking does not vary systematically with age in these cohorts. Moreover, neither the results from the Freiburg monosyllabic word test nor the HSM test affected any response variable in the HA or the CI group.

### 3.3 Speaker bias

In a previous study, a preference in CI users towards the female speaker was found, as quantified by lower listening effort, higher comprehension scores, and preferential neural decoding. With additional data from TH and HA participants, we now investigated if these effects are specific to the CI group. The results are reported in Fig. 5.

**Fig. 5.**
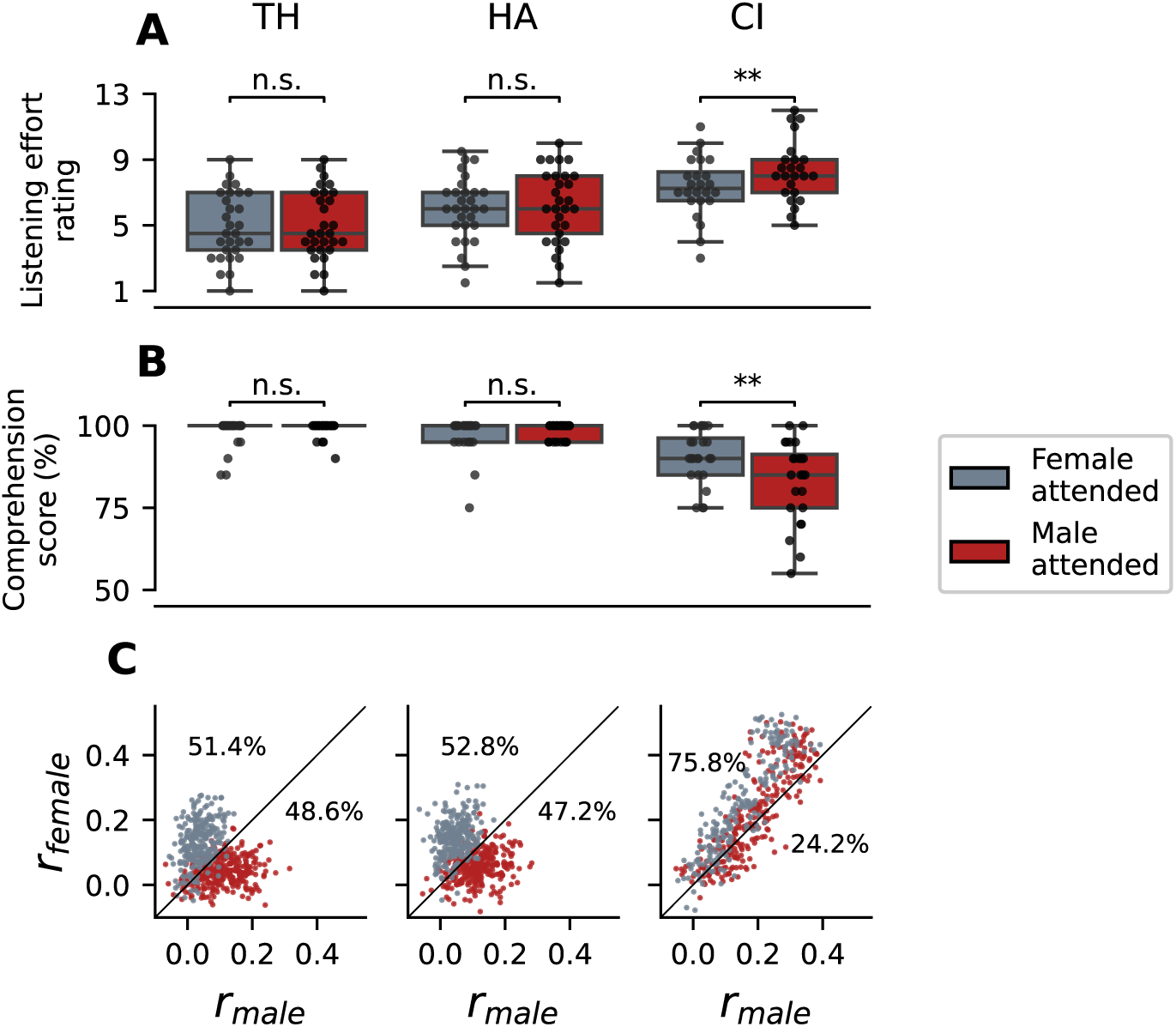
Influence of male vs. female speaker. (**A**) Listening effort ratings were not affected by the speaker identity in the TH and HA groups. However, CI users reported significantly increased listening effort when they were focusing on the male speaker. (**B**) TH and HA participants scored at ceiling for both speakers; no differences are observable. In contrast, CI users answered comprehension questions with higher accuracy when the female speaker was attended. (**C**) Reconstruction scores for the TH and HA group cluster predominantly above the diagonal when the female speaker was attended and below the diagonal otherwise, leading to good separability of classes. In the TH group, 51.4% of all segments appear above the diagonal, indicating that the distribution is almost balanced around the diagonal. With 52.8% of samples above the diagonal, the TH group shows a similarly balanced distribution. Clusters of datapoints in the CI group are not as easily separable. Moreover, 75.8% of samples above the diagonal reveal a substantial bias of the decoder towards the female speaker. Significance notation: ***:*p <* 0.001, **: *p <* 0.01, *: *p <* 0.05, n.s.: *p >* 0.05.

#### 3.3.1 Behavior

Behavioral results in Panels **A** and **B** include data from the single-speaker and competing-speaker conditions. Each data point represents a subject’s average for listening effort of comprehension scores for one speaker. In both cases, multiple testing (within the three cohorts) was compensated for by applying FDR-correction for three comparisons to all p-values.

Panel **A** shows the within-group comparison of listening effort between trials in which the male speaker was attended and those in which the female speaker was attended. As assumptions of normality and homogeneity of variance were met for all groups and sub-groups, we employed paired two-sided t-tests to investigate within-group differences. Within both the TH and HA groups, the median values of listening effort were identical for both speakers: 4.5 and 6.0, respectively. Neither group showed any significant within-group differences, and the effect sizes were negligible (TH: *t*(28) = 0.740, *p* = 0.466, *d* = 0.041; HA: *t*(28) = 0.790, *p* = 0.466, *d* = 0.083). In contrast, the CI group exhibited an increased listening effort for the male speaker (median = 8.0) compared with the female speaker (median = 7.25). This difference was statistically significant (*t*(23) = 3.98, *p* = 0.002), with a medium effect size (*d* = 0.520). The p-values of the paired t-tests were corrected for three comparisons.

We report the within-group comparison of comprehension scores in Panel **B**. As normality was not met for most distributions, we employed non-parametric two-sided Wilcoxon signed-rank tests. Both the TH and HA groups scored at ceiling with median comprehension scores of 100% in both conditions. Consequently, we found no significant within-group differences for either the TH group (*n* = 29; *W* = 3.0, *p* = 0.331, *r*_rb_ = 0.60) or the HA group (*n* = 29; *W* = 18.0, *p* = 0.331, *r*_rb_ = 0.35). Note that the seemingly large effect sizes here are artifacts of the ceiling performance, which drastically reduces the effective sample size to only the few participants who did not score 100%.

In contrast, the CI group (*n* = 24) answered comprehension questions more accurately when the female speaker was attended (median = 90%) than when the male speaker was attended (median = 85%). A two-sided Wilcoxon signed-rank test confirmed that this difference was significant (*W* = 22.0, *p* = 0.006), with a large effect size (*r*_rb_ = *−*0.79). Altogether, speaker-specific behavioral differences were evident only in the CI group.

#### 3.3.2 Neural decoding

To investigate potential biases in neural decoding of the two speakers, we analyzed reconstruction scores from the linear backward model. Rather than averaging across subjects, we show pairs of reconstruction scores for the male and the female speaker (*r_male_*, *r_female_*), each computed on a 60 s segment, to better illustrate the distribution of the data. As before, we used twelve-fold cross-validation, with competing speaker trials serving as test sets. Given an average trial duration of two minutes, we obtained approximately 24 reconstruction scores per participant. Additionally, we calculated the percentage of samples above (i.e. *r_female_ > r_male_*) and below (i.e. *r_female_ < r_male_*) the diagonal, to quantify the balance of the distributions around the diagonal.

Fig. 5 **C** shows similar clustering patterns for the TH and HA groups: data points of the attended female speaker cluster predominantly above the diagonal, while those of the attended male speaker cluster below. This clear separation explains the high decoding accuracies reported for the TH and HA group in Fig. 4 **C**. The distributions are also approximately balanced around the diagonal, with 51.4% of samples in the TH group and 52.8% in the HA group above the diagonal. Consistent with lower decoding accuracy in the CI group, the clustering of reconstruction scores for the male and female speaker is notably less pronounced. Moreover, we find a substantial distribution shift towards the female speaker, quantified by 75.8% of all data points appearing above the diagonal. Together, these findings suggest that the observed preference toward the female speaker is specific to the CI group.

### 3.4 Forward model

As a complementary approach to our primary objective, we also used forward models to investigate the neural encoding of speech. The results generated by forward models for the three cohorts are presented in Fig. 6. The following section includes comparisons of encoding scores between attended, ignored, and null models - the latter computed on circularly shifted versions of the speech envelope. In addition, analyses of model weights and topographies are provided to further characterize neural encoding patterns.

**Fig. 6.**
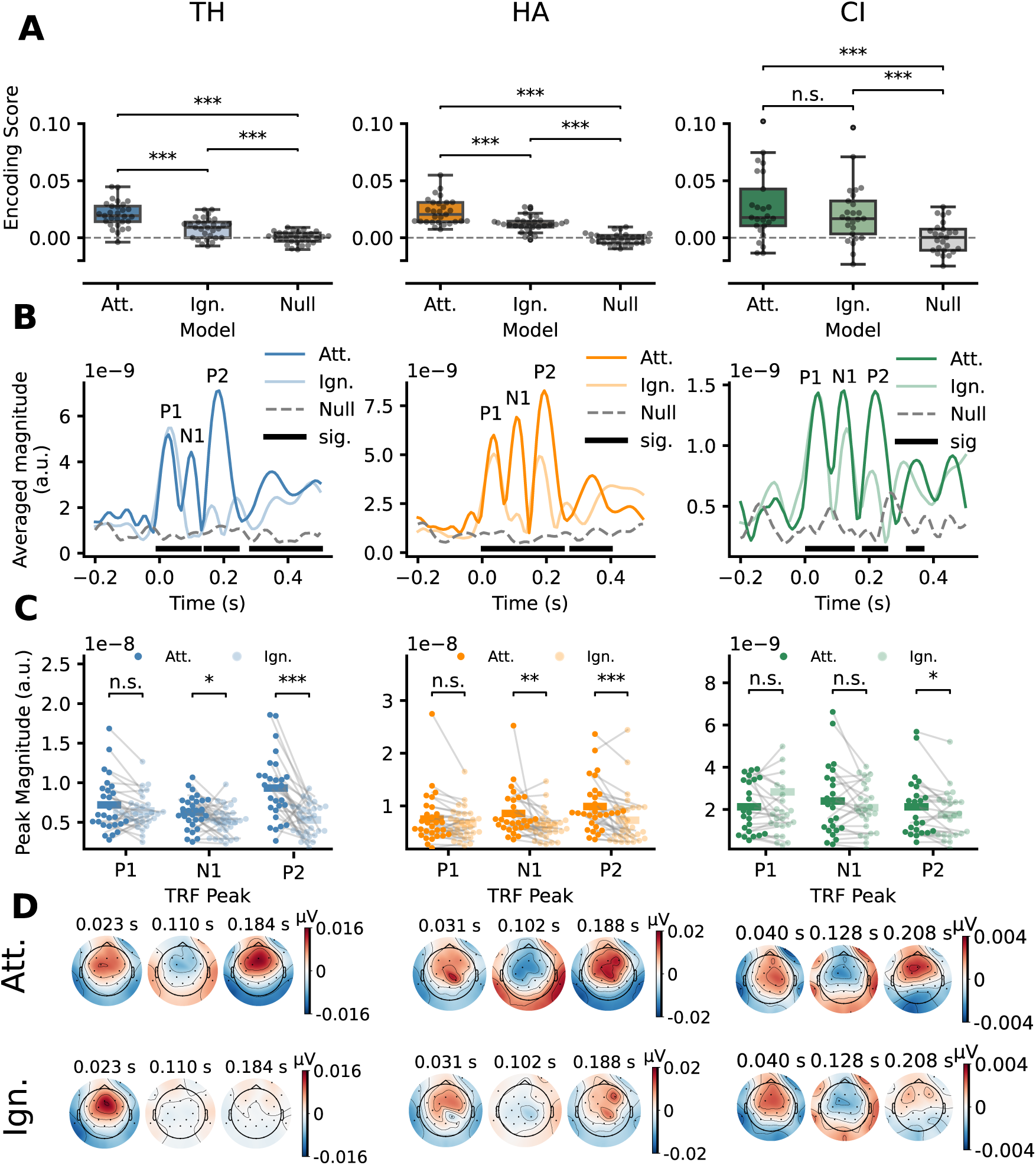
Forward modeling results across the three listener cohorts. Models were computed using the attended, ignored, and a circularly shifted version of the attended speech envelope (null model). (**A**) Encoding scores show a stepwise, significant decrease from attended to ignored to null in the TH and HA groups. In the CI group, attended and ignored scores did not differ significantly, though both exceeded the null model. (**B**) Bold lines below the TRF indicate lags where the attended TRF significantly exceeds the null distribution. All cohorts show three characteristic TRF peaks. Whereas in all groups the *P* 1*_T RF_* peak is unaffected by attention, the *N* 1*_T RF_* is significantly modulated by attention in the TH and HA groups. Similarly, the *P* 2*_T RF_* component is significantly stronger in the attended model across all cohorts. (**C**) Comparison of subjects’ TRF magnitudes that provide statistical evidence for attentional modulation of the TRF peaks. (**D**) Topographic visualization of the TRF weights at the local maxima of the attended TRF prior to averaging and rectification. Fronto-central activation as a response to the attended stimulus is apparent across lags for all groups. In agreement with the analysis of peak magnitudes, ignored topographies reflect weaker responses of *N* 1*_T RF_* for the TH and HA group, as well as weaker responses at *P* 2*_T RF_* across all groups. Significance notation: ***:*p <* 0.001, **: *p <* 0.01, *: *p <* 0.05, n.s.: *p >* 0.05.

#### 3.4.1 Encoding scores

In Fig. 6 **A**, we report the encoding scores, computed using five-fold cross-validation with 20% of every competing speaker trial as a validation set. Within each cohort, comparisons were made between the attended, ignored, and null models. As the assumption of homogeneity of variance between the three models was not met for any of the groups and normality did not hold for attended scores in the HA group, we employed non-parametric significance tests. Within each cohort, we compared the models with a Friedman test, and, if significant, followed up with pairwise comparisons using a two-sided Wilcoxon signed-rank test. The p-values of the non-parametric tests were FDR corrected for three and nine comparisons, respectively.

The TH cohort showed significant differences in encoding scores across models (*χ*^2^(2) = 35.37, *p <* 0.001), with a large effect size (Kendall’s *W* = 0.61). The attended model yielded significantly higher encoding scores than the ignored model (*W* = 20.0, *p <* 0.001), with a very large effect size (*r*_rb_ = 0.91), indicating that, as a result of attention, the neural signal could be encoded more accurately by the attended envelope. Comparisons with the null model further proved that both models performed significantly better than chance, with very large effect sizes observed in both cases (attended vs. null: *W* = 4.0, *p <* 0.001, *r*_rb_ = 0.98; ignored vs. null: *W* = 42.0, *p <* 0.001, *r*_rb_ = 0.81).

The HA group likewise exhibited significant differences among model scores (*χ*^2^(2) = 50.48, *p <* 0.001), with a very large effect size (Kendall’s *W* = 0.87). In line with the findings from the TH cohort, the attended model produced significantly higher encoding scores than the ignored model (*W* = 16.0, *p <* 0.001), again with a very large effect size (*r*_rb_ = 0.93). Moreover, both the attended and ignored models yielded significantly higher scores compared to the null model, showing perfect or near-perfect effect sizes (attended vs. null: *W* = 0.0, *p <* 0.001, *r*_rb_ = 1.0; ignored vs. null: *W* = 2.0, *p <* 0.001, *r*_rb_ = 0.99). This indicates that both the attended and the ignored speech stream contributed to the neural signal, albeit to different extents.

The CI group also showed significant differences in encoding scores across models (*χ*^2^(2) = 11.12, *p* = 0.004), though with a small effect size (Kendall’s *W* = 0.22). Although a tendency towards higher encoding scores of the attended model compared to the ignored model was observable (*median*_att_ = 0.0180 vs. *median*_ign_ = 0.0168), the difference did not reach statistical significance (*W* = 96.0, *p* = 0.075), with a medium effect size (*r*_rb_ = 0.41).

Thus, attentional modulation was not reliably captured by the forward models in this cohort. This highlights limitations of forward models for attention decoding on challenging datasets, such as those from CI users. Backward models, which map from high-dimensional neural activity to a single speech feature, offer a more robust alternative by leveraging differential weighting across EEG channels. Nevertheless, both the attended and ignored models significantly outperformed the null model, showing large effect sizes in both cases (attended vs. null: *W* = 40.0, *p <* 0.001, *r*_rb_ = 0.75; ignored vs. null: *W* = 33.0, *p <* 0.001, *r*_rb_ = 0.80). This indicates that speech-related neural information is present from both speech streams.

Taken together, participants in the TH and HA groups exhibited encoding scores that were clearly modulated by selective attention. The fact that both attended and ignored models consistently outperformed the null model across all cohorts confirms the reliability of the model fits and supports the validity of subsequent analyses based on their weights.

#### 3.4.2 TRFs

We analyzed the TRFs for all three cohorts and for attended versus ignored voices, at different latencies (Fig. 6 **B**). Following standard convention, we identified the first prominent positive peak as *P* 1*_T RF_*, the subsequent negative peak as *N* 1*_T RF_*, and the second positive peak as *P* 2*_T RF_*. All three cohorts exhibited a fourth, smaller peak around 400 ms. As our study focuses on attentional modulation, we will discuss the first peaks in more depth. Because the TRF waveforms were rectified for analysis, their polarity is not apparent in the time-course plots. Instead, the polarity of each component is revealed by its corresponding topographic maps. As the assumption of normality was violated for most distributions, non-parametric two-sided Wilcoxon signed-rank tests were applied. P-values were corrected for multiple comparisons using the FDR method across nine tests.

##### Typical hearing group

The attended TRF displays three distinct peaks with local maxima at 23 ms (*P* 1_TRF_), 110 ms (*N* 1_TRF_), and 184 ms (*P* 2_TRF_), all of which significantly exceed the null distribution. In contrast, the ignored TRF shows only the early *P* 1_TRF_ peak, which is comparable in magnitude to that of the attended condition, while the *N* 1_TRF_ and *P* 2_TRF_ peaks are largely absent. This observation is corroborated by the statistical comparison of individual TRF peak magnitudes shown in Fig. 6 **C**. Here, a two-sided Wilcoxon signed-rank test indicated no significant difference in *P* 1_TRF_ coefficients between attended and ignored conditions (*W* = 139.0, *p* = 0.118), with a medium effect size (*r*_rb_ = 0.36). Moreover, both the attended and ignored TRF topographies exhibit fronto-central positivity of comparable magnitude, though the ignored model displays a slightly sharper, more centrally focused distribution (Fig. 6 **D**). Both later peaks show significantly greater amplitudes in the attended condition, with a medium effect size for the *N* 1_TRF_ (*W* = 108.0, *p* = 0.030, *r*_rb_ = 0.50) and a very large effect size for the *P* 2_TRF_ (*W* = 40.0, *p <* 0.001, *r*_rb_ = 0.81). As shown in Panel **D**, the *N* 1_TRF_ peak is characterized by a diffuse but clearly visible negativity over the fronto-central region, while the *P* 2_TRF_ peak exhibits a sharply defined positivity in the same area. In contrast, the ignored *N* 1_TRF_ and *P* 2_TRF_ topographies are near zero across the entire scalp, highlighting the attentional modulation of later auditory responses. These findings are consistent with previous literature, in which the *P* 1_TRF_ component is primarily attributed to early acoustic encoding that is minimally influenced by attention, while the *N* 1_TRF_ and *P* 2_TRF_ components reflect higher-order processing that is strongly modulated by selective attention.

##### Hearing aid group

The attended TRF exhibits three distinct peaks with local maxima at 31 ms (*P* 1_TRF_), 102 ms (*N* 1_TRF_), and 188 ms (*P* 2_TRF_). The ignored TRF shows a *P* 1_TRF_ peak of comparable magnitude, while the *N* 1_TRF_ is largely absent. The *P* 2_TRF_ peak remains clearly identifiable in the ignored condition but is reduced in magnitude compared to the attended TRF. A comparison of individual TRF magnitudes reveals no significant difference in *P* 1_TRF_ peak amplitudes between conditions (*W* = 139.0, *p* = 0.118), with a small-to-medium effect size (*r*_rb_ = 0.36). In Panel D, the topographies show fronto-central positive activation at the *P* 1_TRF_ peak for both attended and ignored responses. In contrast, the later peaks exhibit significant attentional modulation: both *N* 1_TRF_ and *P* 2_TRF_ are significantly larger in the attended condition, with large effect sizes observed for both peaks (*N* 1_TRF_ attended vs. ignored: *W* = 61.0, *p* = 0.001, *r*_rb_ = 0.71; *P* 2_TRF_ attended vs. ignored: *W* = 56.0, *p <* 0.001, *r*_rb_ = 0.74). The *N* 1_TRF_ peak displays a more pronounced fronto-central negativity in the HA group compared to the TH group, while the ignored TRF remains near zero across the scalp. The attended *P* 2_TRF_ in the HA group is comparable in amplitude to that of the TH group. Notably, unlike the TH group, the HA group’s ignored TRF shows distinct fronto-central activity, although it remains significantly weaker than the attended response. Overall, the HA group demonstrates a TRF pattern comparable to that of the TH group. However, attentional modulation in the HA cohort is somewhat stronger at the *N* 1_TRF_ peak, as reflected by the larger effect size and more distinct topographies. The attentional modulation at the *P* 2_TRF_ peak remains highly significant but slightly attenuated compared to the TH group.

##### Cochlear implant group

As shown in Fig. 6 **B**, the TRF peak magnitudes of CI participants are markedly reduced compared to those observed in the TH and HA cohorts. This attenuation is likely a consequence of the CI-specific artifact rejection procedure, which diminishes overall EEG signal power. Despite the lower amplitudes, however, three distinct peaks are identifiable at 40 ms (*P* 1_TRF_), 128 ms (*N* 1_TRF_), and 208 ms (*P* 2_TRF_). As in the other two cohorts, the *P* 1_TRF_ magnitudes are nearly identical between attended and ignored conditions, a finding supported by the subject-level comparison (*W* = 115.0, *p* = 0.330), which showed a small effect size (*r*_rb_ = 0.23). Likewise, topographies in Panel **D** show similar negative activation in the fronto-central area of the attended and ignored model. The *N* 1_TRF_ peak in the attended condition appears slightly larger than in the ignored condition, though the difference is less pronounced than in the TH and HA groups and does not survive statistical comparison (*W* = 107.0, *p* = 0.258), with a small effect size (*r*_rb_ = 0.29) as shown in Panel **C**. Correspondingly, the topographies in Panel **D** reveal similar spatial patterns for both conditions, with slightly stronger activation in the attended TRF. In contrast, the *P* 2_TRF_ component shows a clear difference between conditions, with the ignored TRF exhibiting a significantly reduced magnitude, as shown in Panel **C** (*W* = 61.0, *p* = 0.0215), with a large effect size (*r*_rb_ = 0.59). In line with this analysis, the topographies in Panel **D** show the greatest distinction at the *P* 2_TRF_ peak: the attended TRF displays strong fronto-central positivity, while the ignored TRF shows minimal activation across the scalp. Overall, these findings indicate that in the CI group, attentional modulation emerges during later stages of neural processing.

## 4 Discussion

Our study conducted a comprehensive investigation of cortical speech tracking across age-matched users of HAs, CIs, as well as typically hearing (TH) participants. Moreover, it used a realistic free-field paradigm with participants using their own clinically fitted devices. Through complementary encoding and decoding analyses as well as a speaker-specific decoding analysis, our work offers one of the most thorough assessments to date of how users of assistive hearing technology neurally track speech in complex auditory scenes. This closes a gap in current research, as cortical responses to competing speech have been studied a) in isolation for either hearing aid (HA) (Decruy et al., 2020; Fuglsang et al., 2020; Petersen et al., 2017) or cochlear implant (CI) users (Nogueira et al., 2019; Paul et al., 2020), but did not perform a direct comparative analysis of these two key assistive technologies. Furthermore, we introduce more ecologically valid scenarios (e.g., use of own clinically fitted devices, free-field stimulation). Also, while previous studies have documented a behavioral preference for female speakers among CI users (El Boghdady et al., 2019; Gaudrain et al., 2023), the neural basis for this phenomenon has remained an open question until now.

### 4.1 Behavioral responses

#### Emerging differences in the competing-speaker scenario

In our study, participants answered comprehension questions and rated listening effort. In the single-speaker scenario, all three cohorts performed at ceiling in the comprehension task with no group differences. For the listening effort, the CI group demonstrated slightly higher effort than the other two groups. However, this difference was not statistically significant either. These findings are consistent with previous research showing that both HA and CI users typically achieve satisfactory speech understanding in quiet environments (Litovsky et al., 2017; Zeng et al., 2008). After introducing a competing speaker, however, substantial group differences emerged. Although subjective listening effort increased for all groups, including the TH group, there was now a significant increase in listening effort from the TH to the HA group, and further from the HA to the CI group. Moreover, comprehension scores were significantly lower in the CI group compared to both other cohorts. The majority of participants in the TH and HA groups still performed at ceiling. Although TH and HA did not differ significantly in their comprehension scores, the fact that more participants in the HA group (9 individuals) than in the TH group (4 individuals) scored below 100% suggests a tendency towards increased comprehension difficulties in the HA group.

#### The role of spatial release from masking

We hypothesize that the observed group differences in behavior for the competing-speaker scenario stem from different levels of spatial release from masking, i.e., the benefit of spatial separation of target and distractor in a complex auditory scene. TH participants are typically able to localize sound in the horizontal plane through interaural time differences (ITDs) and interaural level differences (ILDs). The different spatial locations of the speakers in our experiment, separated by 60^◦^ in the horizontal plane, probably allowed our TH participants to separate the target from the distractor very well, thus allowing better focus on the target. The hearing loss in HA and CI users presumably meant that they could benefit less from spatial release from masking, despite using hearing assistive devices (Litovsky et al., 2017). Although bilateral hearing aid users can make use of both ITDs and ILDs to localize sound in the horizontal plane (Litovsky et al., 2017), it has been shown that spatial release from masking (SRM) in bilateral HA users is around 8 dB less than in TH individuals (Marrone et al., 2008). This effect may account for the increased listening effort observed in this group compared to TH participants.

#### Limitations of CI stimulation

CI processors primarily transmit envelope information, resulting in the loss of the speech signal’s temporal fine structure. Additionally, the processors for each ear typically operate independently, which can lead to unsynchronized stimulation. Both of these factors hinder sound localization and speech understanding in complex acoustic environments (Litovsky et al., 2017). Unlike TH individuals, CI users rely predominantly on ILDs for sound localization, as their ITD thresholds are substantially higher than those of TH listeners (Aronoff et al., 2010; Bernstein and Trahiotis, 2002). Bilateral CI users, hence exhibit only modest SRM, typically around 2-4 dB, and the performance gap compared to TH listeners is especially pronounced when the distractor is one competing speaker, as in our paradigm (Loizou et al., 2009). Overall, the absence of temporal fine structure in CI stimulation, elevated ITD thresholds and limited SRM benefits together can account for the lowest performance observed in CI users in the competing-speaker paradigm, and aligns with the known difficulties this population group has in multi-speaker environments.

### 4.2 Neural encoding and decoding

We assessed the neural tracking of amplitude fluctuations in speech through encoding and decoding models of the speech envelope. We quantified the performance of the decoding models through the reconstruction score of the speech envelope as well as through the accuracy of attention decoding, which was based on the decoding models. The encoding models were investigated through their TRFs as well as the resulting encoding scores. The TRFs exhibited three significant peaks at latencies around 20 ms, 100 ms, and 200 ms for which attentional effects were quantified. A further, smaller peak occurred around 400 ms. Neural tracking at these longer latencies was presumably related to the N400 component of event-related potentials and has previously been shown to be related to semantic processing, reflecting, for instance, semantic dissimilarity between words as well as surprisal in a sequence of words (Broderick et al., 2018; Hahne et al., 2024; Weissbart et al., 2020).

#### Hearing-aid users

##### Cortical tracking comparable to that of typical hearing participants

One objective of the neural analysis was to determine if differences in cortical speech tracking could explain the increased listening effort and lower comprehension in competing-speech situations observed in the HA population. Our findings, however, suggest that cortical speech tracking between HA users and TH individuals is largely convergent. Thus, increased listening effort and decreased comprehension could not be explained by differences in cortical speech tracking. Specifically, reconstruction scores for attended speech were highly comparable, and decoding accuracies were indistinguishable. Although a slight, but statistically insignificant, increase was noted in the reconstruction of ignored speech for the HA cohort relative to the TH group, the overall pattern points towards robust and similar neural processing in the two cohorts. The similarity of neural tracking in the HA and the TH groups was further supported by the TRF analysis, where both groups were found to exhibit significant attentional modulation of encoding scores. Again, the encoding scores of the ignored envelope were slightly increased for the HA group compared to the TH group. Moreover, both cohorts exhibited significant attentional modulation of both the *N* 1*_T RF_* and *P* 2*_T RF_* peaks. In contrast, prior work (Decruy et al., 2020; Fuglsang et al., 2020) found increased overall envelope tracking and attentional modulation in HA users as compared to TH subjects, as quantified by classification accuracy. A crucial difference in study design likely explains this discrepancy: Whereas these prior studies provided linearly amplified stimuli over headphones, our study analyzed a more ecological scenario, where participants utilized their standard hearing aids and settings in a free-field acoustic setting.

##### Critical role of hearing aids in normalizing speech tracking

The advanced signal processing in modern hearing aids thus appears to effectively restore the acoustic signal and to normalize the listening experience for HA users. This restoration may reduce the need for the compensatory neural gain that is otherwise required to track a degraded signal, thereby causing similar cortical speech tracking as in typical hearing participants. This interpretation is supported by our finding that within the TH (hearing loss *≤* 25 dB), greater hearing loss was a significant predictor of increased neural tracking of the ignored speaker. This result is consistent with previous work showing that non-target speech was better predicted by an attended decoder for hearing-impaired individuals (Fuglsang et al., 2020) and that hearing loss was associated with increased tracking of ignored speech (Petersen et al., 2017). The presence of a hearing loss effect within the TH group, but the absence of an inter-group difference between the TH and the HA group further emphasized the critical role of hearing aids in normalizing cortical speech tracking.

##### Persistent Listening Effort in Hearing Aid Users

Our analysis of cortical speech tracking did not explain why hearing aid users continue to struggle in multi-talker situations, as evidenced by their higher listening effort in our study and previous reports (Harkins and Tucker, 2007; Miles et al., 2022; Shinn-Cunningham and Best, 2008). A potential explanation may lie in other stages of auditory processing, such as subcortical responses at higher frequencies, which may reveal underlying deficits in temporal processing that are not compensated for by current hearing aid technology (Forte et al., 2017; Maddox and Lee, 2018; Schüller et al., 2023; Skoe and Kraus, 2010). Also, as mentioned before, bilateral HA users profit less from spatial release from masking, which would also explain an increase in listening effort despite good cortical speech tracking (Marrone et al., 2008).

#### CI users

##### Reduced attentional modulation

Overall, the CI group showed significantly reduced cortical attentional modulation, i.e., the ability to focus on the target and ignore the distractor. This became apparent in the analysis of both the decoding and the encoding model. In contrast to the HA individuals, the CI group was characterized by substantially increased reconstruction scores from the backward model for both attended and ignored speech, a result attributable to stimulation artifacts. However, the difference between attended and ignored reconstruction scores, despite being robust, was smaller in the CI group than in the other groups, resulting in lower decoding accuracies. This pattern of high overall reconstruction scores combined with weak attentional modulation is highly consistent with prior findings (Nogueira et al., 2019), where stimuli were streamed directly to the CIs. Our study thus demonstrates that this phenomenon persists in a more ecological free-field listening environment. The weaker attentional modulation of the cortical speech tracking was even more pronounced in the forward model’s encoding scores. Despite overall successful neural prediction of both attended and ignored speech, as evidenced by the respective encoding scores significantly exceeding the null model, encoding scores did not differ significantly between attended and ignored speech. In line with prior work (Paul et al., 2020), significant attentional modulation for the CI users was only evident at the *P* 2*_T RF_* peak, while the earlier *N* 1*_T RF_* only exhibited a tendency towards being more pronounced in the attended condition. Our findings highlight the profound deficit in attentional modulation in the CI users as a potential cause for their speech-in-noise comprehension difficulties. Methodologically, the relative success of the backward model highlights its robustness in decoding attention from noisy or artifact-heavy data compared to the forward model, that required artifact removal for interpretable TRFs.

##### Artifact removal

Although persistent artifact contamination is always a concern with CI data, the results presented in Fig. 6 support robust artifact rejection due to two key findings. Firstly, since CI artifacts are highly correlated and time-locked to speech signal, a crucial indicator of CI artifacts would be a pronounced peak in the TRF at a latency of around 0 ms. However, our TRF analysis did not reveal any substantial response at a latency of 0 ms, suggesting effective removal of CI-related artifacts (Fig. 6B).

Secondly, the high correlation between CI artifacts and the speech envelope would artificially inflate the encoding scores of the forward model beyond what is expected for neural responses. Yet, after artifact removal, our encoding scores of the attended speaker (Fig. 6 B) drop to a level comparable to that of the other populations (medians: TH: 0.019, HA: 0.020, CI: 0.017). The encoding scores of the ignored speaker remain higher than in the other two groups (medians: TH: 0.009, HA: 0.012 CI: 0.016). However, we think this nonetheless reflects the neural responses only, since CI artifacts should affect both target and distractor scores equally.

##### Correlation with comprehension scores

While a positive correlation between neural tracking and speech comprehension is expected (Etard and Reichenbach, 2019), our results differed across listener groups due to differences in behavioral performance. For the TH and HA groups, comprehension scores were at ceiling even for the competing speaker scenario, precluding a meaningful analysis of this relationship. Conversely, the CI group displayed a wider distribution of comprehension scores, which revealed a significant positive relationship between decoding accuracy and comprehension. This finding directly corroborates prior work (Nogueira and Dolhopiatenko, 2022; Verschueren et al., 2019) and highlights that for CI users, the strength of neural tracking is a key determinant of their ability to understand speech in noise.

### 4.3 Speaker bias

#### Neural evidence for female-voice advantage in CI users

Our use of a male and a female talker revealed a significant speaker-specific bias in the CI group. Behaviorally, CI users reported lower listening effort and achieved higher comprehension scores when attending to the female voice. This behavioral advantage was mirrored in the neural data, as a speaker-specific decoder analysis showed that the female talker was decoded more often as the attentional target than the male one. Crucially, this effect was absent in both the TH and HA groups under identical testing conditions, indicating that the bias is not an experimental confound but rather a characteristic of CI processing.

#### Relationship with the fundamental frequency and vocal tract length

To understand the origin of this speaker bias, it helps to consider the key acoustic cues that differentiate the talkers. The most critical of these are the fundamental frequency (*F* 0), which determines perceived pitch, and the vocal tract length (VTL), which correlates with the speaker’s perceived size (Fitch and Giedd, 1999). Typically, female speakers are characterized by a higher F0 and a shorter VTL. The stimuli in our experiment reflected this pattern, with the female voice having a higher f0 (164.5 Hz vs. 110.9 Hz) and a shorter VTL (14.2 cm vs. 16.7 cm)

Our finding of a female voice preference is consistent with a complementary set of findings in the literature. For instance, (Gaudrain et al., 2023) showed that when target and masker were presented at the same level—a condition analogous to our study—CI users benefit from acoustic differences when the masker has more male characteristics. This complements prior work (El Boghdady et al., 2019), where it was observed that performance worsened as the masker became more female-like, an effect they attributed to an unfavorable target-to-masker ratio developing in the CI electrodogram.

Taken together, these studies suggest a consistent pattern: CI performance is optimized when the target voice is female and the masker is male. Our study strongly supports this interpretation by providing clear behavioral evidence of a female target preference and, for the first time, also showing its neural basis. This reinforces the critical need to investigate stimulation strategies that can accommodate or correct for this perceptual bias, a line of research advocated for instance by El Boghdady et al. (2021).

### 4.4 Limitations

#### The interpretation of our findings should take into account the following limitations

First, results for the CI group cannot necessarily be generalized to the entire CI user population, as participants were required to achieve a minimum level of speech perception in quiet and to successfully complete the experimental protocol. This introduces a sampling bias that underrepresents CI users with poor outcomes and reduces the observed variability relative to the full clinical population. However, the applied inclusion criterion (*≥* 60% on the Freiburg monosyllabic word test at 65 dB SPL) corresponds to a common performance range for postlingually deafened adult CI users assessed with this test rather than an exceptionally high-performing subgroup (Czurda et al., 2024; Franke-Trieger et al., 2023; Hoppe et al., 2019, 2023). Notably, even within this group behavioral and neural scores in the competing speaker condition remained lower in the CI group than in the TH or HA group, underscoring the particular challenge that multi-talker listening situations pose even for CI users with typical or above-average speech perception in quiet.

Second, for the TH and HA groups, comprehension scores were at ceiling, which precluded a meaningful analysis of the relationship between neural tracking and behavioral performance due to a lack of variance in the behavioral data. Additionally, while the use of personal hearing aid settings was central to our objective of studying cortical tracking in an ecological setting, it introduced variability in signal processing across participants.

Third, the EEG data from CI users was inherently confounded by significant electrical artifacts stemming from the CI. Although CI artifact removal is a necessary step for the TRF data visualization and evaluation, the differently treated forward model data make direct comparisons across groups more challenging. However, within-subject effects—particularly attentional modulation—remain robust and interpretable as they rely on relative changes within the same recording conditions.

A fourth limitation of the current study is the lack of data on specific binaural CI performance. This would inform us to what degree CI users are able to use spatial release from masking, which is a likely factor contributing to the decreased comprehension scores and attentional modulation observed in the CI group. Future studies would benefit from measuring metrics such as ILD and ITD sensitivity in this cohort to assess the interaction between bilateral CI performance and selective neural tracking.

One should also consider that our analysis had to be run mostly with non-parametric tests. This means that due to the reduced power of these tests, we may miss or underestimate some effects.

### 4.5 Conclusion

This study is the first to compare neural tracking of cochlear-implant users, hearing-aid users, and typically hearing participants and also the first to do so in a challenging free-field competing-speaker paradigm with participants using their own clinically fitted devices.

For cochlear implant users, our results show a profound impairment in the neural segregation of attended and ignored speech at the level of cortical speech tracking. This was also reflected by the significantly increased behavioral difficulties in the competing speaker condition, where CI users exhibited the highest listening effort and the lowest speech comprehension. Furthermore, we are the first to demonstrate a neural basis for the perceptual bias of CI users favoring female voices in a multi-talker setting.

In contrast, we found that hearing-aid users exhibit largely normal-like cortical tracking when using their own devices. This suggests that modern hearing aids largely normalize the cortical tracking of speech envelopes. Persistent difficulties may stem from uncompensated temporal processing deficits, which could be identified by probing subcortical responses to high-frequency information in future research, and from a reduced spatial release from masking.

The highly divergent outcomes of the two hearing-impaired groups highlight the need to understand neural processing in realistic listening conditions to successfully develop next-generation stimulation strategies tailored to specific, device-dependent deficits.

## Acknowledgements

We sincerely thank all participants for their willingness to take part in this study, without whom this work would not have been possible. We are grateful to Maxi Wollenberger, who carried out a substantial part of the data acquisition with great dedication, and to Domenic Schulze for generously sharing his expertise on hearing-aid users and providing guidance on appropriate testing procedures. We also thank Mike Thornton for engaging and insightful discussions on the methodology of forward and backward modelling.

## Declaration of generative AI and AI-assisted technologies in the writing process

During the preparation of this work the authors used ChatGPT and Google AI Studio in order to improve readability and language. After using this tool, the authors reviewed and edited the content as needed and take full responsibility for the content of the published article.

## Data availability

The data recorded for this study is publicly available. We published three distinct datasets:

- The data of the Cochlear Implant group are available under https://doi.org/10.5281/zenodo.17952844
- The data of the Hearing Aided group are available under https://doi.org/10.5281/zenodo.17927767
- The data of the Typically Hearing group are available under https://doi.org/10.5281/zenodo.17952231

Source-code for data analysis is publicly available on https://github.com/Constantin-Jehn/aad-neuroimage.git.

## A Appendix: Impact of years of CI-use on cortical tracking

A potential factor impacting the attentional modulation of cortical speech tracking is CI experience. To study this putative predictor, we fitted two additional model for decoding accuracy in CI-users.

Firstly, we fitted a model that added the mean duration of implant use, indicating the average years of implant use across ears (*duration_mean_*). This model then read:

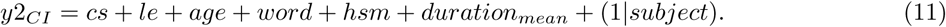

Secondly, we fitted a model that added the maximal duration of implant use (i.e. the years of implant since the first implantation) (*duration_mean_*) to the model. This model read:

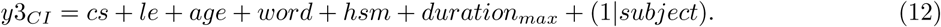

The results are summarised in Table 3. Both models yielded a decreased model fit (y2: log-likelihood = *−*193.0, y3: log-likelihood = *−*193.75) compared to the initial model (log-likelihood = *−*192.75). Consistently, both predictors did not show significant predictive power, as evidenced by high p-values (y2: 0.56, y3:0.92)

**Table 3:** Comparison of extended Linear Mixed Effects Models with CI experience predictors.

| Model | Added Predictor | Log-likelihood | Predictor $\beta$ (95% CI) | p-value |
| --- | --- | --- | --- | --- |
| y2 | Mean duration of implant use | $-193.50$ | $-0.093 [-0.406, 0.220]$ | 0.560 |
| y3 | Maximal duration of implant use | $-197.75$ | $-0.016 [-0.321, 0.291]$ | 0.920 |

## B Appendix: Individual CI data

**Table 4:** Demographics and CI details of CI users at the individual level.

| ID | Age (yr) | Sex | CI Exp. right (yr) | CI Exp. left (yr) | Implant right | Implant left | Processor right | Processor left | Coding Str. right | Coding Str. left | Monosyllable right (65 dB %) | Monosyllable left (65 dB %) | HSM 65/60 dB | Etiology / Clinical Course |
| --- | --- | --- | --- | --- | --- | --- | --- | --- | --- | --- | --- | --- | --- | --- |
| 102 | 62 | M | 9 | 6 | HiRes90K Mid-Scala | HiRes90K Midscala | Naida CI M90 | Naida CI Q90 | HiRes Optima-S | HiRes-S incl. Fidelity 120 | 60 | 60 | n/a | Noise induced (progressive) |
| 103 | 45 | M | 6 | 10 | CI422(SRA) | CI522 | CP910 | CP910 | ACE | ACE | 75 | 80 | n/a | Unknown |
| 104 | 56 | F | 16 | 10 | CI422RE(CA) | CI422(SRA) | CP910 | CP1000 | ACE | ACE | 90 | 60 | 11 | Unknown |
| 105 | 58 | F | 16 | 20 | Pulsar | C40+ | SONNET | SONNET | FS4 | FS4 | 70 | 55 | 13 | Unknown |
| 106 | 62 | M | 9 | 8 | CI422(SRA) | CI522 | CP1150 | CP1150 | ACE | ACE | 80 | 85 | 61 | Idiopathic SNHL, Progressive |
| 107 | 57 | M | 10 | 4 | Mi1000 FLEX 28 | Mi1200 FLEX 28 | SONNET | SONNET | FS4 | FS4 | 85 | 90 | 0 | Unknown (progressive) |
| 108 | 56 | M | 4 | 7 | Mi1200 FLEX 28 | Mi1250 FLEX 28 | SONNET | SONNET | FS4 | FS4 | 80 | 90 | 53 | Idiopathic SNHL, Progressive |
| 109 | 36 | F | 8 | 5 | Mi1200 FLEX 28 | Mi1200 FLEX 28 | SONNET | SONNET | FS4 | FS4 | 80 | 90 | 58 | Unknown (progressive) |
| 110 | 30 | M | 28 | 23 | C40 | C40 | SONNET 2 | SONNET 2 | HDCIS | HDCIS | 85 | 80 | 24 | Congenital |
| 111 | 26 | F | 15 | 4 | Sonata | CI522 | SONNET 2 | CP1000 | FSP | ACE | 50 | 90 | 8 | Congenital (progressive) |
| 112 | 69 | F | 12 | 11 | Sonata | Mi1000 Std | SONNET | SONNET | FS4 | FS4 | 90 | 80 | 25 | Unknown (progressive) |
| 113 | 81 | M | 6 | 7 | HiRes90K Mid-Scala | HiRes90K Mid-Scala | Naida CI M90 | Naida CI M90 | HiRes Optima-S | HiRes Optima-S | 75 | 75 | 16 | Unknown (progressive) |
| 114 | 41 | F | 13 | 16 | CI512 | CI24RE | CP910 | CP910 | ACE | ACE | 90 | 80 | 9 | Congenital (progressive) |
| 116 | 65 | M | 14 | 4 | CI24RE(CA) | CI522 | CP1000 | CP1000 | ACE | ACE | 90 | 70 | 25 | Otosclerosis |
| 118 | 27 | F | 17 | 25 | CI24RE(CA) | CI24M | CP910 | CP910 | ACE | ACE | 40 | 85 | 0 | Congenital |
| 119 | 37 | M | 6 | 4 | CI522 | CI522 | CP910 | CP1000 | ACE | ACE | 80 | 90 | 36 | Unknown (progressive) |
| 120 | 30 | F | 15 | 2 | CI24RE(CA) | CI622 | CP1150 | CP1150 | ACE | ACE | 85 | 75 | 23 | Congenital |
| 121 | 61 | F | 17 | 22 | Pulsar | C40+ | SONNET | SONNET | FSP | FSP | 80 | 80 | 53 | Otosclerosis |
| 122 | 67 | F | 5 | 17 | CI512 | CI24RE | CP1110 | CP1110 | ACE | ACE | 85 | 75 | 32 | Meningitis |
| 123 | 28 | F | 25 | 12 | Pulsar | Mi1200 FLEX 28 | SONNET 2 | SONNET 2 | FS4 | FS4 | 85 | 20 | 17 | Congenital |
| 125 | 25 | F | 12 | 12 | CI512 | CI512 | CP1000 | CP1000 | ACE | ACE | 100 | 95 | 52 | Idiopathic SNHL |
| 127 | 65 | M | 7 | 2 | HiRes90K Mid-Scala | HiResUltra 3D SlimJ | Naida CI M90 | Naida CI M90 | HiRes Optima-S | HiRes Optima-S | 85 | 80 | 8 | Unknown (progressive) |
| 128 | 56 | F | 5 | 2 | CI522 | CI622 | CP1000 | CP1000 | ACE | ACE | 80 | 80 | 82 | Unknown (progressive) |
| 130 | 75 | F | 24 | 13 | CI24M | CI512 | CP1000 | CP1000 | ACE | ACE | 75 | 80 | 3 | Unknown (progressive) |

